# Neuro-glial lipid imbalance in a *Drosophila* model of Amyotrophic Lateral Sclerosis 8

**DOI:** 10.64898/2026.06.06.730557

**Authors:** Lovleen Garg, Kriti Chaplot, Gowhith Kuppili, Shweta Tendulkar, Jomon Joseph, Siddhesh Kamat, Girish Ratnaparkhi

## Abstract

Membrane Contact sites (MCS) have emerged as physiologically relevant zones that coordinate inter-organelle communication and cellular function. VAPB, an ER-resident MCS tethering protein, plays a central role in regulating MCSs through its numerous protein interactors, thereby influencing cellular homeostasis. A pathogenic missense *VAPB^P56S^* mutation causes familial Amyotrophic Lateral Sclerosis 8 (ALS8) in humans, with progressive degeneration of motor neurons. The precise mechanisms underlying the motor neurodegeneration remain poorly understood.

In this study, we examine lipid imbalance in the brain of a *Drosophila* model of ALS8 (*VAPB^P58S^)*. Specifically, we find that lipid homeostasis is disrupted in an age-dependent manner. Strikingly, cholesterol esters and sphingolipids show an age-dependent increase, while cholesterol shows a decrease. Intriguingly, from a cellular perspective, despite the accumulation of triacylglycerols (TAGs) in the brains of *VAPB^P58S^* animals, the increased neutral lipid species do not correlate with lipid droplets (LDs), which are fewer in density and smaller in size. Lipid imbalance and progressive motor dysfunction in *VAPB^P58S^* animals can be reversed by expressing *VAPB^WT^*, suggesting a relationship between VAPB activity and lipid flux.

To uncover VAPB’s role in lipid homeostasis, we modulate *VAPB* activity in neurons and glia to dissect out tissue-specific roles. We find that both cell types contribute to lipid homeostasis in differential ways. In glia, LD flux is strongly dependent on VAPB activity, a dependence further recapitulated in cultured human cell lines, suggesting evolutionary conservation of the regulatory mechanism.

Thus, we hypothesise that lipid dysregulation constitutes a critical pathogenic feature of ALS8, with the *VAPB^P56S^* allele disrupting lipid homeostasis in the neuro-glial axis.

**Summary Statement:** The ER-membrane tethering protein VAPB regulates lipid homeostasis

## Introduction

Amyotrophic Lateral Sclerosis (ALS) is an adult late-onset progressive neurodegenerative disease characterised by the loss of both upper and lower motor neurons (van Es *et al*. 2017). Neuronal degeneration leads to muscle weakness, loss of voluntary movement, and thus paralysis in the patient (Genge *et al*. 2024). Early symptoms of ALS are muscle cramps, spasticity and localized muscle weakness in the limbs and/or neck. As the disease progresses, patients suffer slurred speech, difficulty in swallowing food, sialorrhea, shortness of breath and respiratory failure (Liguori *et al*. 2021). Most of the patients die within 2-5 years after diagnosis. The exact cellular mechanisms that lead to ALS remain unknown and are poorly understood, with no effective cure (Alsultan *et al*. 2016; Genge *et al*. 2024). Most ALS cases are sporadic (sALS), with the underlying cause unknown, while ∼10% are familial (fALS), transmitted across generations (Ajroud-Driss and Siddique 2015). Although fALS cases are quite low compared to sALS, their studies have been instrumental in identifying genetic causes and disease mechanisms (VAN Damme *et al*. 2017). To date, more than 30 genes have been associated with disease, including 18 definitive loci. These loci are classified into various functional categories like RNA metabolism, mitochondrial function, proteostasis, ER stress, cytoskeleton dynamics and protein quality control (Teuling *et al*. 2007; Shi *et al*. 2010; Dupuis *et al*. 2011; Robelin and Gonzalez de Aguilar 2014; Costa and de Carvalho 2016; Xu *et al*. 2016). Given the molecular heterogeneity, multiple mechanisms are likely to converge to produce common clinical and pathological phenotypes characteristic of ALS (Layalle *et al*. 2021).

Emerging evidence from the past decade highlights lipid metabolism as an important factor in ALS pathogenesis (Desport *et al*. 2001; Desport *et al*. 2006; Dupuis *et al*. 2008; Dupuis *et al*. 2011; Costa and de Carvalho 2016; Gonzalez de Aguilar 2019). As many alterations are observed in patients with ALS, it is crucial to study the underlying causes (Agrawal *et al*. 2022). ALS loci like *TARDBP* and *SOD1* have been shown to play important roles in lipid metabolism (Iguchi *et al*. 2012; Chaves-Filho *et al*. 2019). In the healthy brain, oxidative and metabolic stress is generally low; however, hypermetabolism in ALS increases energy demand and reduces nutrient availability. Moreover, malnutrition and symptoms of loss of appetite have been observed in ALS patients (Browne *et al*. 2006; Sol *et al*. 2021; Tracey *et al*. 2021). A significant decrease in membrane fluidity in the brain and spinal cord has also been observed as ALS progresses in mouse models, possibly due to lipid peroxidation (Miana-Mena *et al*. 2011). Alterations in polyunsaturated/saturated lipid ratio can decrease membrane fluidity and promote lipid-lipid and lipid-protein cross-linking. These changes can thus affect motor neuron function in ALS. Such disruptions have been proposed as a potential mechanism underlying pathology in ALS, further emphasising the role of lipids in its pathogenesis (Chaves-Filho *et al*. 2019).

In a Brazilian family of 28 affected members distributed across four generations, an autosomal dominant missense mutation, Proline 56 to Serine, in *VAMP-associated protein B* (*VAPB)* was identified as the 8^th^ locus of ALS (*ALS8*, (Nishimura *et al*. 2004)). *ALS8* patients generally show symptoms of progressive muscle weakness, cramps, autonomic dysfunction and behavioural impairments (Mcbenedict *et al*. 2025). ALS8 subtypes not only differ in symptoms but also in disease onset and progression (Chen *et al*. 2010; Sanhueza *et al*. 2015; Murage *et al*. 2024). Other ALS mutations identified in the *VAPB locus* are P56H, T46I, A145V and V234I (Kors *et al*. 2022a). The *VAPB^P56S^* mutation, modelled as *VAP^P58S^* in *Drosophila*, leads to the misfolding of the mutant protein, disrupts cellular functions such as membrane trafficking and the ER stress response, and activates the unfolded protein response (UPR) (Teuling *et al*. 2007; Moustaqim-Barrette *et al*. 2014; Tendulkar *et al*. 2022; Thulasidharan *et al*. 2024).

VAPB is a Type IV integral ER membrane protein with three domains major sperm protein (MSP), coiled-coil domain (CCD) and transmembrane domain (TMD) (James and Kehlenbach 2021; Neefjes and Cabukusta 2021). The MSP domain is present at the N-terminus of the protein and faces towards the cytosol and is known to interact with other proteins to define membrane-membrane contact sites (MCS) (Kamemura and Chihara 2019; Bashar *et al*. 2025)). The VAPB MSP domain can be cleaved, secreted, and perform extracellular functions, as in *C. elegans* and *Drosophila* (Han et al. 2012; Kamemura and Chihara 2019). The CCD is a site for interaction with other VAP family proteins and SNARES (Wyles and Ridgway 2004; Murphy and Levine 2016). The C-terminal TMD is inserted into the ER membrane (James and Kehlenbach 2021). VAPB is ubiquitously expressed in mammals and conserved across species. It interacts with a variety of proteins to perform cellular functions such as organelle tethering, lipid transfer, calcium homeostasis, autophagy, proteostasis and UPR (Baron *et al*. 2014; Costello *et al*. 2017; Gomez-suaga *et al*. 2017; Kirmiz *et al*. 2018; Zhao *et al*. 2018; Kamemura *et al*. 2021; Zein-sabatto *et al*. 2021; Kors *et al*. 2022b; Kors *et al*. 2022c; Kalarikkal *et al*. 2024; Ji *et al*. 2025). Several MSP binding partners contain the FFAT motif (two phenylalanines in an acidic tract), which is a part of the canonical sequence (EFFDA-E) flanked by acidic residues;

Binding partners also interact with VAPB using non-canonical sequences (Mikitova and Levine 2012; Murphy and Levine 2016). To study the role of *VAPB* in the context of ALS8, various cellular and animal models have been generated, including patient-derived iPSCs, mutant VAP-expressing *S. cerevisiae, M. musculus*, *R. rattus*, *C. elegans,* and *D. melanogaster*(Teuling *et al*. 2007; Ratnaparkhi *et al*. 2008; Han *et al*. 2012; Tran *et al*. 2012; Deivasigamani *et al*. 2014; Moustaqim-barrette *et al*. 2014; Sanhueza *et al*. 2014; Tokutake *et al*. 2015; Chaplot 2019; Tendulkar *et al*. 2022; Stump *et al*. 2023; Murage *et al*. 2024; Thulasidharan *et al*. 2024). Each model has its own advantages and disadvantages. Across various studies, ER stress, UPR, mitochondrial function, and inflammation are among the common factors shown to contribute to the progression of ALS (Teuling *et al*. 2007; Moustaqim-barrette *et al*. 2014; Tendulkar *et al*. 2022; Stump *et al*. 2023). However, the role of lipid metabolism in ALS8 has been poorly investigated.

Our laboratory works on a fly ortholog of VAPB (*VAP33A*, *CG5014; VAP* henceforth), and we have used it to generate an overexpression model of *VAP^WT^*/*VAP^P58S^* for a genomic screen to define a genetic sub-network that includes *VAP* (Deivasigamani *et al*. 2014). Further, we have uncovered *SOD1* and *Tor* as genetic interactors of *VAP* that modulate *VAP* aggregation (Chaplot 2019) via reactive oxygen species (ROS). Knockdown of these genes also led to an increased amount of oxidised lipids due to higher ROS levels. Then we used the ALS8 fly genomic line, in which the mutant gene *VAP^P58S^* is expressed under the native *VAP33A* promoter, as described (Moustaqim-barrette *et al*. 2014; Tendulkar *et al*. 2022). We showed that Caspar/FAF1, a *VAP* interactor, was involved in regulating glia-mediated inflammation via the NF-κB pathway in the context of ALS8 (Tendulkar *et al*. 2022) and that age-dependent inflammation is a characteristic feature of the *VAP^P58S^* mutant (Kulkarni *et al*. 2026). In the current study, we sought to understand the role of *VAPB* in maintaining lipid homeostasis in the brain, using *Drosophila* as a model organism. With a combination of genetics and biochemistry, we investigated the effect of *VAP*^P58S^ on *Drosophila* brain lipid levels. Based on lipid profiling results, we explored VAP function in glial LD homeostasis. Our study uncovers the role of *VAP* in regulating glial LD homeostasis, which is perturbed in the *VAP^P58S^ Drosophila* model. This disruption in lipid regulation highlights a previously underexplored function of VAPB in the nervous system and underscores the importance of lipid homeostasis in ALS8 pathogenesis.

## Results

### Age-dependent LC-MS-based lipid profiling of the adult *Drosophila* brains in *VAP^P58S^*

A *Drosophila* model for *ALS8* was generated by the Tsuda lab (Moustaqim-barrette *et al*. 2014). In this model, a first chromosome VAP null (*ΔVAP*) line, which is larval/pupal lethal, was rescued either by transgenic *VAP^WT^*or by a *VAP^P58S^* genomic insert on the 3^rd^ chromosome. Both alleles were expressed by a native/genomic *VAP* promoter, and the respective lines, as utilised by our laboratory (Tendulkar *et al*. 2022; Thulasidharan *et al*. 2024), are hereafter referred to as *ΔVAP;gVAP^P58S^* or *ΔVAP;gVAP^WT^*. Here, *ΔVAP* (on X) is rescued by a *single copy* of the genomic VAP (*gVAP*) allele, inserted on the 3^rd^ Chromosome. Both alleles could rescue lethality, with the crucial difference that, post-eclosion, the *ΔVAP;gVAP^P58S^* line showed a shorter lifespan (median survival age 25 vs 50 days for wild-type) and age-dependent deterioration of motor (climbing) function (Fig. 1A) (Moustaqim-barrette *et al*. 2014; Tendulkar *et al*. 2022), whereas the *ΔVAP;gVAP^WT^* was on par with wild-type. 1 to 5-day-old *ΔVAP;gVAP^P58S^* animals showed normal climbing, as measured by the ‘Startle-Induced Negative Geotaxis, (SING)’ assay, but older flies (5 to 30-day-old) showed a progressive loss of motor function with age, with animals losing all ability to move or climb after 25 days (Fig. 1A). Flies with a double copy of g*VAP^P58S^* (*ΔVAP;gVAP^P58S^/ gVAP^P58S^*), have a higher density of VAP aggregates, progress through larval and pupal stages, eclose normally, but are short-lived, and their progressive loss of motor function is accelerated when compared to a single copy of *gVAP^P58S^* (Moustaqim-barrette *et al*. 2014; Tendulkar *et al*. 2022; Thulasidharan *et al*. 2024).

**Figure 1:**
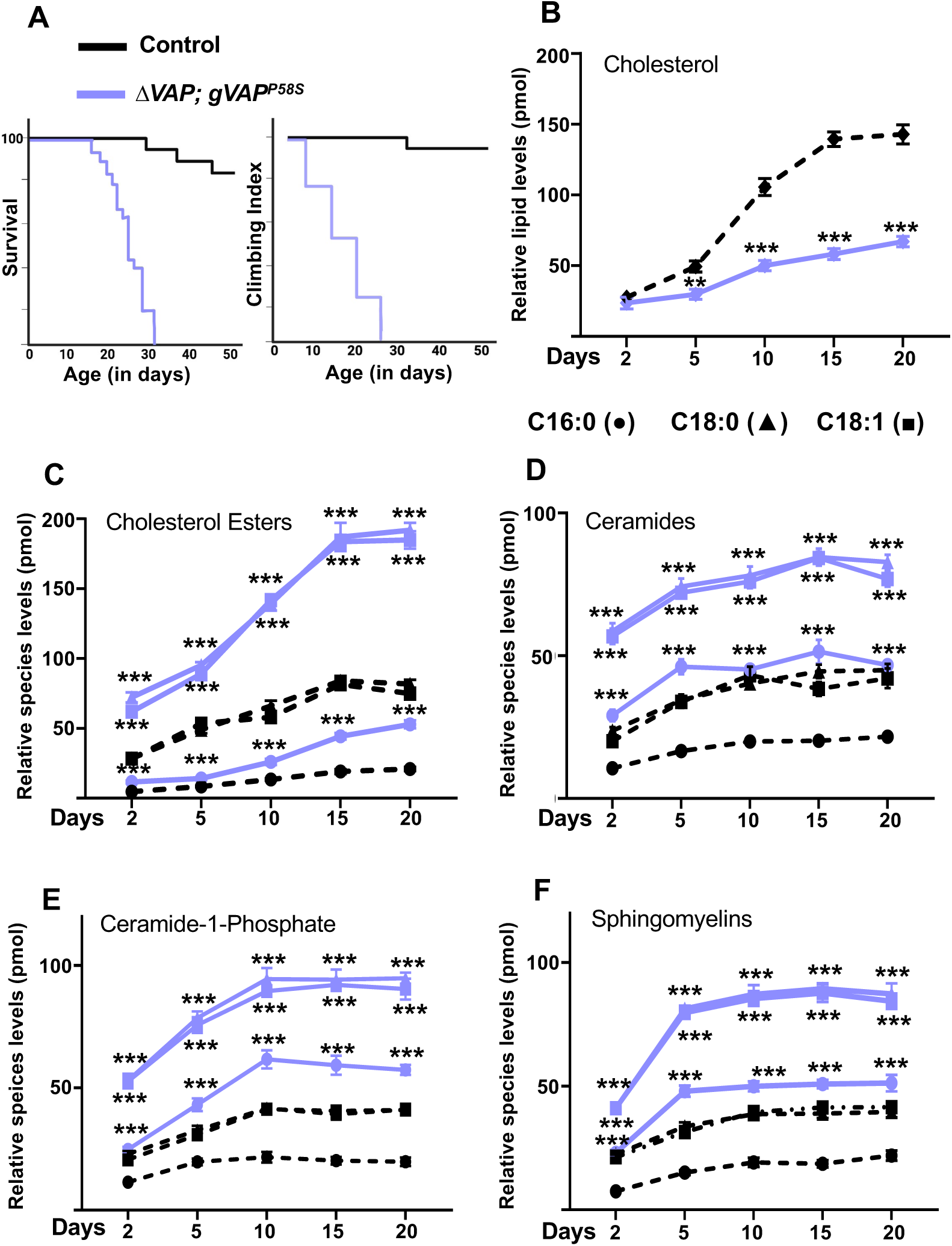
Age-dependent lipid profiling of the *Drosophila* adult brain. **(A)** Schematic representation of the survival curve and climbing index of control (black) and *ΔVAP;gVAP^P58S^* (light purple) males. **(B)** Cholesterol levels in the adult brain in *ΔVAP;gVAP^P58S^* (light purple) are at par with controls (black) at day-2. Intriguingly, cholesterol levels did not rise steeply in *ΔVAP;gVAP^P58S^* brains, with age, unlike controls **(C)** In contrast, CEs are elevated at day-2 in *gVAP^P58S^* (light purple) and their concentration in the adult brain increases with age, as compared to controls-(black). **(D-F)** Ceramides (panel D), Ceramide-1-Phosphate (panel E) and Sphingomyelins (panel F) show significant age dependent increase in *ΔVAP;gVAP^P58S^* compared to wild-type. Individual lipid species are indicated as C16:0(●), C18:0(▴) and C18:1(▪). Relative lipid species levels are plotted on Y-axis with age (in days) on X-axis. Data is presented as mean±SEM. Statistical tests were performed using multiple unpaired t-tests comparing genotypes at each time point for individual lipid species, followed by the FDR approach of Benjamin and Hochberg for multiple-comparison correction. p values are reported as *(<0.05), **(<0.01) and ***(<0.001). N=5 biological replicates, n=2 age-matched adult brains for each genotype.

Being interested in age-dependent changes in the brain lipid profile, we performed targeted lipidomics on adult brains from both *ΔVAP;gVAP^P58S^* and wild-type males on the 2^nd^, 5^th^, 10^th^, 15^th,^ and 20^th^ day post-eclosion (Fig. 1 B-F). We find that a subset of lipids changes with age in the wild-type (Black symbols/lines; Fig 1B-F), showing a mild to significant increase. *ΔVAP; gVAP^P58S^* brains (light violet symbols/lines; Fig 1B-F) show significant age-dependent deviation when compared to controls.

Flies are cholesterol auxotrophs, with cholesterol obtained from the diet; therefore, it is intriguing that *ΔVAP; gVAP^P58S^* brains do not show a normal age-dependent increase in cholesterol (Fig. 1B), with levels being ∼2.5 fold lower in 20-day animals. This suggests that either cholesterol is not reaching the brain or is being rapidly converted into another lipid species, in comparison with control flies. The abnormally high levels of cholesterol esters (CEs) in the brain (Fig. 1C), with C16:0, C18:0, and C18:1 species increasing with age and being 4-fold higher at day-20, suggest that in *ΔVAP;gVAP^P58S^*, that feeding is normal and possibly cholesterol conversion to CEs is not reversed after reaching the brain. A similar trend is seen for *ΔVAP; gVAP^P58S^* with ceramides, C1P and sphingomyelins, with 2-fold increase, on average, with lipid levels stabilizing by day-10 (Fig. 1D-F).

### Adding a copy of *VAP^WT^* to g*VAP^P58S^* reverses lipid dysregulation

Both, the shortened lifespan and progressive decrease in motor function in *ΔVAP; gVAP^P58S^* males can be rescued by adding a single copy of *gVAP^WT^* (*ΔVAP; gVAP^P58S^/gVAP^WT^*) (Moustaqim-barrette *et al*. 2014; Thulasidharan *et al*. 2024). Also, adding a single copy of *gVAP^WT^*supports the clearance of aggregates in the adult brain (Thulasidharan *et al*. 2024) with increasing age, accompanied by a concomitant reduction in ER stress (Thulasidharan *et al*. 2024). Based on this ability of VAP^WT^ in reversing detrimental effects of VAP^P58S^, we performed a targeted LC-MS based lipid profiling on brain lysates of *ΔVAP;gVAP^P58S^*, *ΔVAP;gVAP^WT^* and *ΔVAP; gVAP^P58S^/gVAP^WT^* 15-day-old males (Fig. 2). Cholesterol levels, which are lower in *ΔVAP;gVAP^P58S^* go back to normal in the brains of 15-day-old *ΔVAP; gVAP^P58S^/gVAP^WT^*. CE’s, which are high in *ΔVAP;gVAP^P58S^*, are normal in *ΔVAP; gVAP^P58S^/gVAP^WT^*, as are C1P, ceramides and sphingomyelins. Intriguingly, Sphingosine and S1P levels are similar across all genotypes evaluated (wild-type, mutant or rescue). Phospholipids and lysophospholipids also do not change across genotypes except for Phosphatidylserine (C34:0) (Suppl. Fig. 1, 2). Neutral lipids, such as TAGs were significantly higher in the 15-day-old *ΔVAP; gVAP^P58S^* brain (Fig. 2G), with levels approaching normal levels after adding a copy of *VAP^WT^*(*ΔVAP; gVAP^P58S^/gVAP^WT^*).

**Figure 2:**
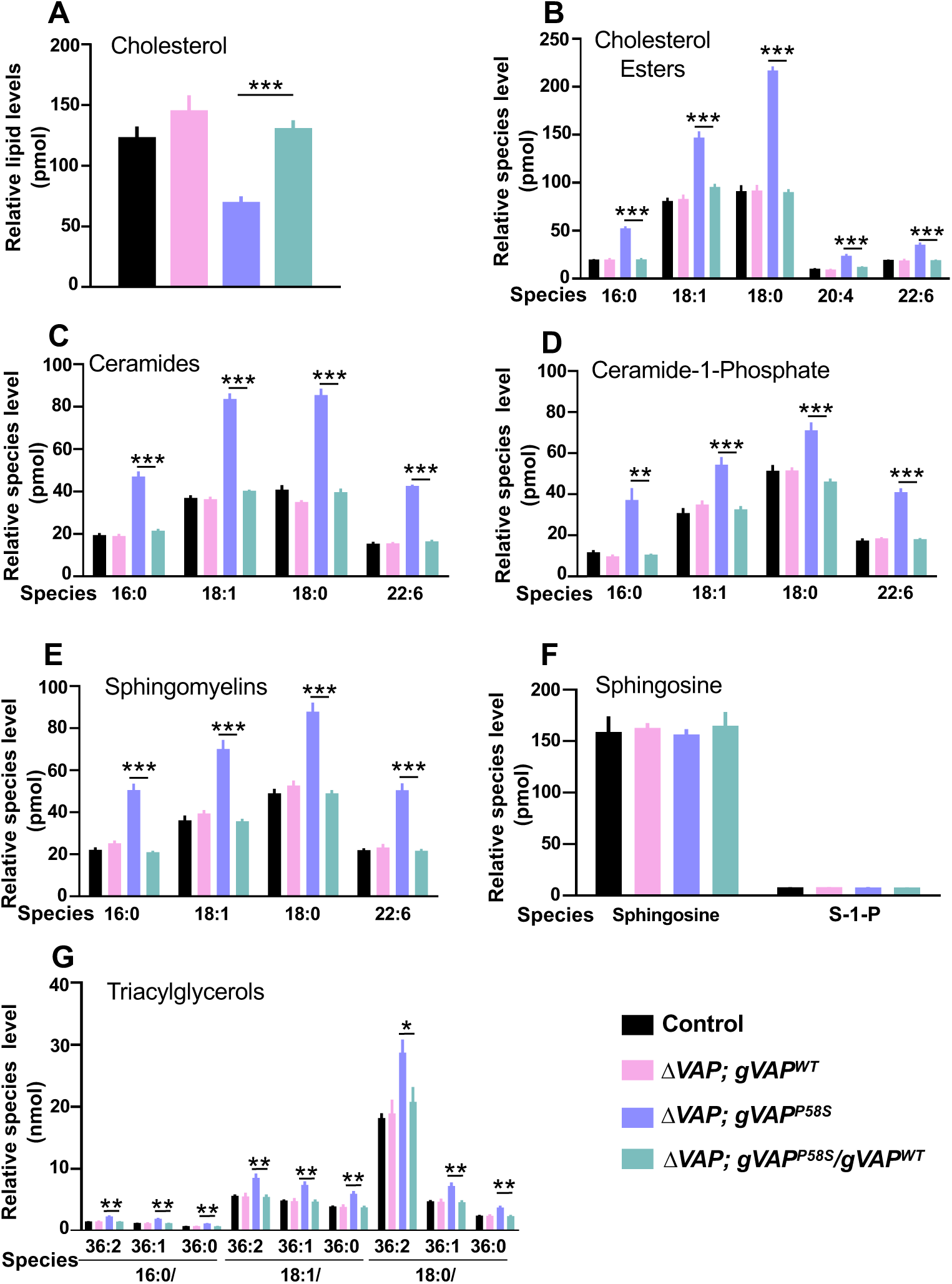
A single copy of *VAP^WT^* restores adult brain lipid levels in the 15-day-old *ALS8* flies. Lipid levels return to control/normal levels upon addition of g*VAP^WT^* in *ΔVAP;gVAP^P58S^* background, for cholesterol **(A)**, cholesterol esters **(B)**, ceramides **(C)**, ceramide-1-phosphate **(D)**, sphingomyelins **(E)** and triacylglycerols **(G)**. No significant changes were detected in sphingosine and sphingosine-1-phosphate in control and mutant **(F)**. Levels of all these lipids remain equivalent between control, *ΔVAP, gVAP^WT^*, and *ΔVAP, gVAP^P58S^/gVAP^WT^*. Relative lipid/species levels (mean±SEM) are plotted on the Y axis with lipid species indicated on the X axis. A multiple unpaired t-test was used to compare means within the same lipid species across different genotypes, followed by the FDR approach of Benjamin and Hochberg for multiple-comparison correction with p values reported as *(<0.05), **(<0.01) and ***(<0.001). N=5 biological replicates, n=2 adult brains per genotype.

Surprisingly, lipid species which act as substrates for the formation of TAGs, such as FFA, MAGs and DAGs, did not change in the 15-day-old *VAP^P58S^* male brains at all (Suppl. Fig. 3A-C). Thus, expression of a single allelic dose of *VAP^WT^* rescued not only behavioural traits, such as lifespan and motor function (Moustaqim-barrette *et al*. 2014; Tendulkar *et al*. 2022; Thulasidharan *et al*. 2024), but also the lipid imbalance observed in the *ΔVAP;gVAP^P58S^* brain. Based on these interesting results, especially the high TAG levels, we examined the density and size of LDs within the brain in an age-dependent manner. This allowed us to explore cellular aspects of lipid imbalance in the ageing brain via imaging. The interest in LDs stemmed from the knowledge that LDs are storage depots for TAGs and CE’s and are increasingly believed to be active metabolic zones in the cell (Bohnert and Schrul 2024; Mathiowetz and Olzmann 2024; Henne *et al*. 2025).

### Age-dependent Lipid Droplet distribution in the *ALS8* model of *Drosophila*

CEs and TAGs are the core components of LDs. This led us to stain and quantify LDs using BODIPY (493/503) following the protocols described by (Garg *et al*. 2026). For LD quantitation, we chose the suboesophageal ganglion (SOG) as our region of interest (ROI), as described in Materials and Methods.

Wild-type animals showed a stable density of LDs in the adult brain, which did not change with age (Fig. 3A-D, day-5 to day-20), when quantified (Fig. 3E, black bars). LD size, however, increased with 20-day-old flies having higher number of larger LDs as compared to day-5 (Fig. 3F, black dots). In contrast, the average LD density decreased (Fig. 3A’-D’ & Fig. 3A’’-D’’; Fig. 3E) in both *ΔVAP; gVAP^P58S^* and *ΔVAP; gVAP^P58S^/gVAP^P58S^*, ranging from a 25-30% drop on day-5 vs a 50% drop in density on day-20 (as compared to wild-type). Unlike wild-type, the average LD size in both *ΔVAP; gVAP^P58S^* and *ΔVAP; gVAP^P58S^/gVAP^P58S^* remained constant (Fig. 3F) with age. This suggested that the VAP^P58S^ variant could sustain LD formation, but in these animals, LD density was lower with fewer large-sized LDs.

**Figure 3:**
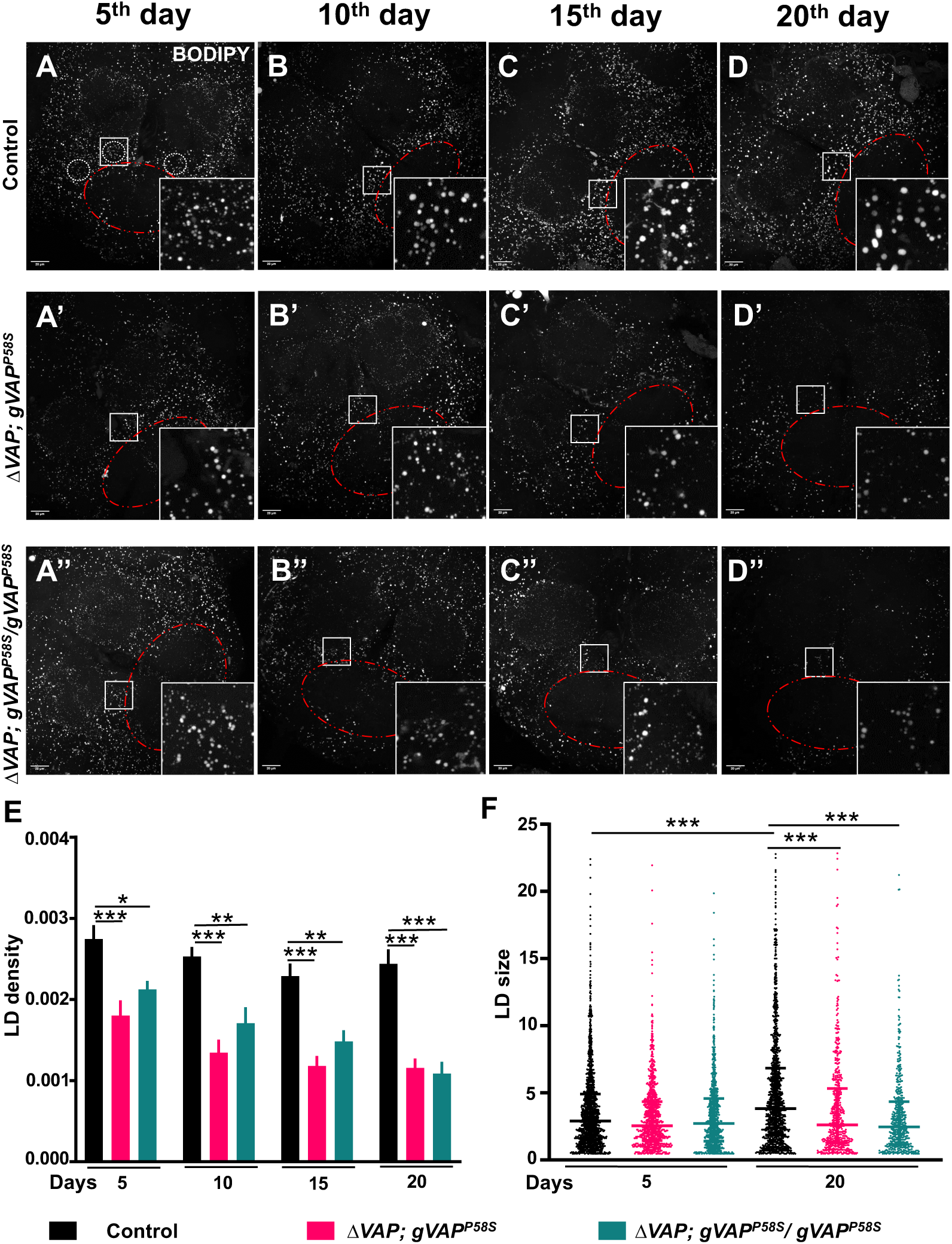
Age-dependent perturbation of lipid droplet homeostasis in the adult brain of *VAP^P58S^*. **(A-D, A’-D’, A’’-D”)** Representative maximum intensity projections (MIPs, 40% Brightness) of BODIPY (493/503) stained lipid droplets in the *Drosophila* adult brain central body (0.75X zoom) with expanded inset on the lower right corner. The scale bar is 20 µm. (A-A”) LD distribution at 5^th^ day for *ΔVAP; gVAP^P58S^* (A’) and Δ*VAP;gVAP^P58S^*/*gVAP^P58S^* (A”) compared to control (A). (B-B”), (C-C”) and (D-D”) show LD distribution at 10^th^, 15^th^ and 20^th^ day respectively. SOG region is selected for the analysis of lipid droplets depicted by red outline. Regions of interest (ROIs) are drawn in control (A) around SOG region selected for LD quantitation. Two ROIs drawn are beneath the zoomed inset (A). **(E)** Quantitation of the LD density in SOG region plotted as mean±SEM of LD numbers per unit volume (µm^3^). A 25-30% drop in LD density is observed on the 5^th^ day in the *ALS8* mutants for both *ΔVAP/Y ;; gVAP^P58S^* and *ΔVAP/Y ;; gVAP^P58S^*/*gVAP^P58S^*. Two-way ANOVA was performed as a statistical test for mean comparisons between genotypes, followed by Tukey’s test for multiple comparison correction with p values reported as *(<0.05), **(<0.01) and ***(<0.001). **(F)** Scattered dot plot for LD size (volume, µm^3^) in SOG region presented with median and interquartile range. LD size increases with age in the wild-type control but not in *ALS8* brains. Kruskal-Wallis test was performed for mean rank comparison of different genotypes, followed by Dunn’s test for multiple comparison correction N=6 adult brains per genotype, n= 5 ROIs analyzed for each brain.

### Adding a copy of *VAP^WT^* to *VAP^P58S^* normalises LD density but not size

Earlier, we found complete rescue of lipid levels in *ΔVAP; gVAP^P58S^* brains after the introduction of a single copy of *VAP^WT^*. Thus, we tested if LDs could also be rescued, both in terms of density and size, in *ΔVAP; gVAP^P58S^*/ *gVAP^WT^* animals. Wild-type animals followed the same pattern (Fig. 4A-C) as seen earlier, with LD density remaining stable (Fig. 4D) from 5-15 days, and average LD size increasing (Fig. 4F) on day-15. The brain of *ΔVAP; gVAP^WT^* animals were at par with controls in terms of LD density and size (Suppl. Fig 4). Here, to maintain an equal dose of *VAP*, we compared *ΔVAP; gVAP^P58S^/ gVAP^P58S^* animals to *ΔVAP; gVAP^P58S^/ gVAP^WT^*. We find an expected drop of LD density by ∼30% (Fig. 4D) in *ΔVAP; gVAP^P58S^/ gVAP^P58S^* (Fig. 4 A’-C’) brains, as also the absence of larger LDs (size >20µm^3^) (Fig. 4E). In brains of *ΔVAP; gVAP^P58S^/ gVAP^WT^* animals (Fig. 4 A’’-C’’), LD density, but not LD size, is rescued to wild-type levels (Fig. 4D,E), with average volumes being lower and a marked absence of LDs of volume >20µm^3^. This suggests that *gVAP^P58S^* inhibits the age-dependent increase in large LDs (> 20 µm^3^), even in the presence of *gVAP^WT^*. This conclusion is supported by experiments in later sections that investigate the effects of *VAP* loss-of-function (*lof*) and gain-of-function (*gof*).

**Figure 4:**
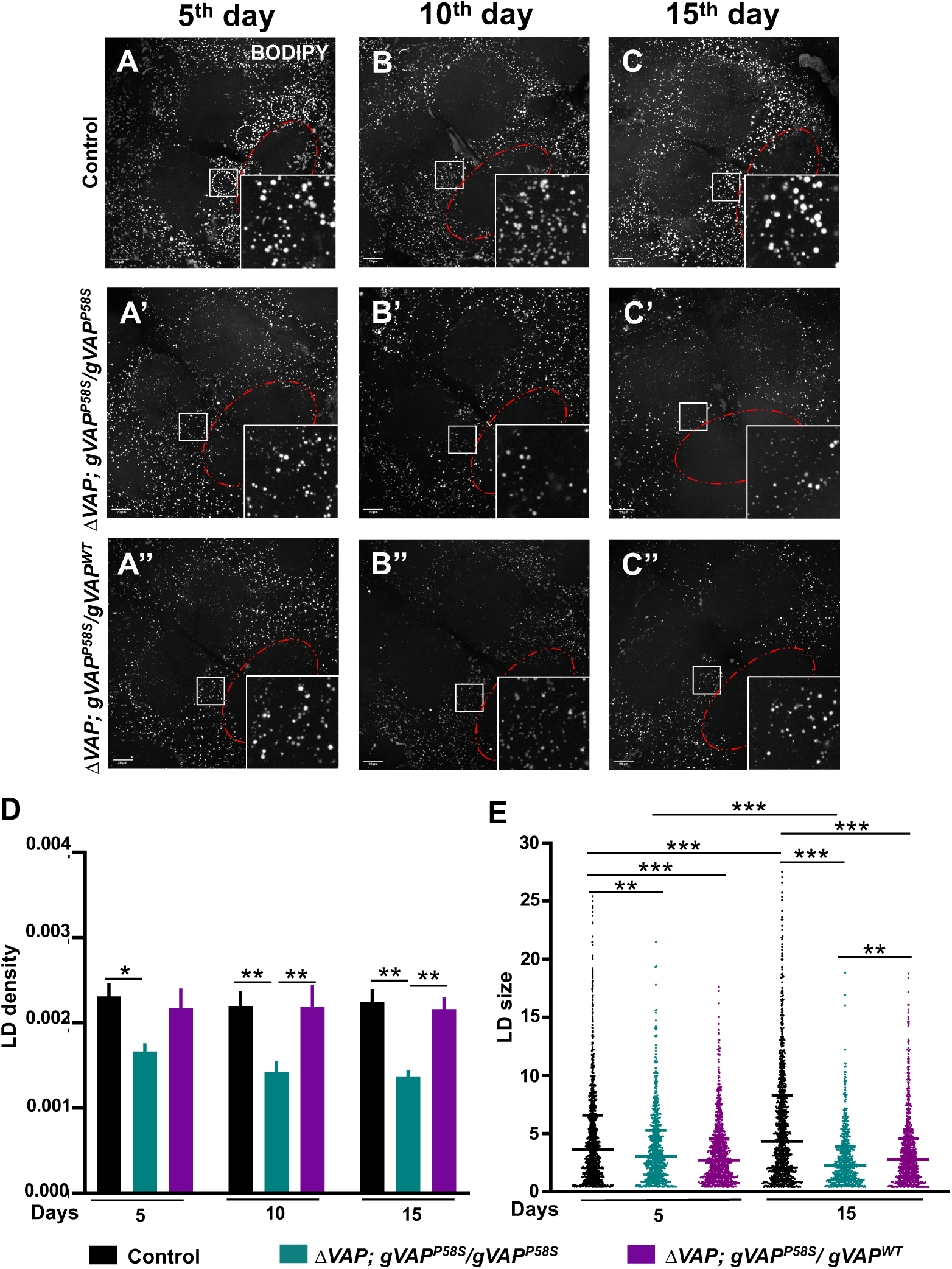
A single copy of *VAP^WT^* restores LD density in *VAP^P58S^* flies. **(A-C”)** Representative MIPs of BODIPY-stained LDs in adult brain’s central body (0.80X zoom) with zoomed inset shown on the lower right corner. Scale bar is shown as 20 µm. **(A-A”)** LD distribution at day-5 for wild-type control (A), *ΔVAP; gVAP^P58S^/gVAP^P58S^* **(A’)** and *ΔVAP;gVAP^P58S^*/*gVAP^WT^* **(A”)**. Similarly, **(B-B”)** and **(C-C”)** represent the distribution for 10^th^ and 15^th^ days. The SOG region is selected for LD analysis, as depicted by a red outline. ROIs are represented around the SOG region in the control **(A)** used for LD quantitation. **(D)** Quantitation for LD density (number of droplets per µm^3^) plotted as mean±SEM values on the Y axis against age (days) on the X axis. LD density is significantly rescued on days 10 and 15 in animals that also express *VAP^WT^*. Two-way ANOVA was performed for statistical test between genotypes, followed by Tukey’s test for multiple comparison correction. **(E)** Scattered dot plot for LD size (volume, µm^3^) is plotted with size on the Y axis, including median with interquartile range, against age (days) on the X axis. The availability of *VAP^WT^* does not rescue LD size in the *ALS8/VAP^P58S^* mutant, but LD size is significantly increased compared to the mutant on the 15^th^ day. Kruskal-Wallis test was used for mean rank comparison between genotypes, followed by Dunn’s test for multiple comparison correction. p values are reported as *(<0.05), **(<0.01) and ***(<0.001). N=5 adult brains per genotype, n= 5 ROIs analyzed for each brain.

### Glial *VAP* modulates brain LD density and size

In previous sections, we found that age-dependent LD homeostasis is perturbed in the brains of *ΔVAP; gVAP^P58S^* animals. Furthermore, LD density can be rescued by the addition of a copy of *gVAP^WT^*, but the largest LDs are not rescued. Since LDs in the brain are primarily found in glial cells (Kis *et al*. 2015; Welte 2015; Girard *et al*. 2020). This suggests that VAP function in glia may contribute to LD homeostasis. To test this hypothesis, we modulated VAP levels by knockdown (*KD*) of *VAP* transcripts via RNA interference (RNAi) and by overexpressing (*OE*) *VAP^WT^* or *VAP^P58S^* in independent experiments.

First, we knocked down *VAP* in glial cells (*repo-Gal4>VAP^RNAi^*) and examined the LD distribution in the adult brain for 5-day and 20-day aged flies (Fig. 5 A-B; Suppl. Fig 5 A-B). LD density on day 5 was marginally higher, whereas on day 20 it was significantly higher (Fig. 5A, B & E) than that of controls. As seen earlier, in wild-type brains, LD size increases with age, with LDs >15 µm^3^ emerging in 20-day brains, and this trend was similar in both control and *VAP* RNAi (Fig. 5B). This suggests that the VAP^WT^ function is important for maintaining LD equilibria in glial cells, with reduced (hypomorphic) *VAP^WT^* in glia leading to an accumulation of LDs in the adult brain.

**Figure 5:**
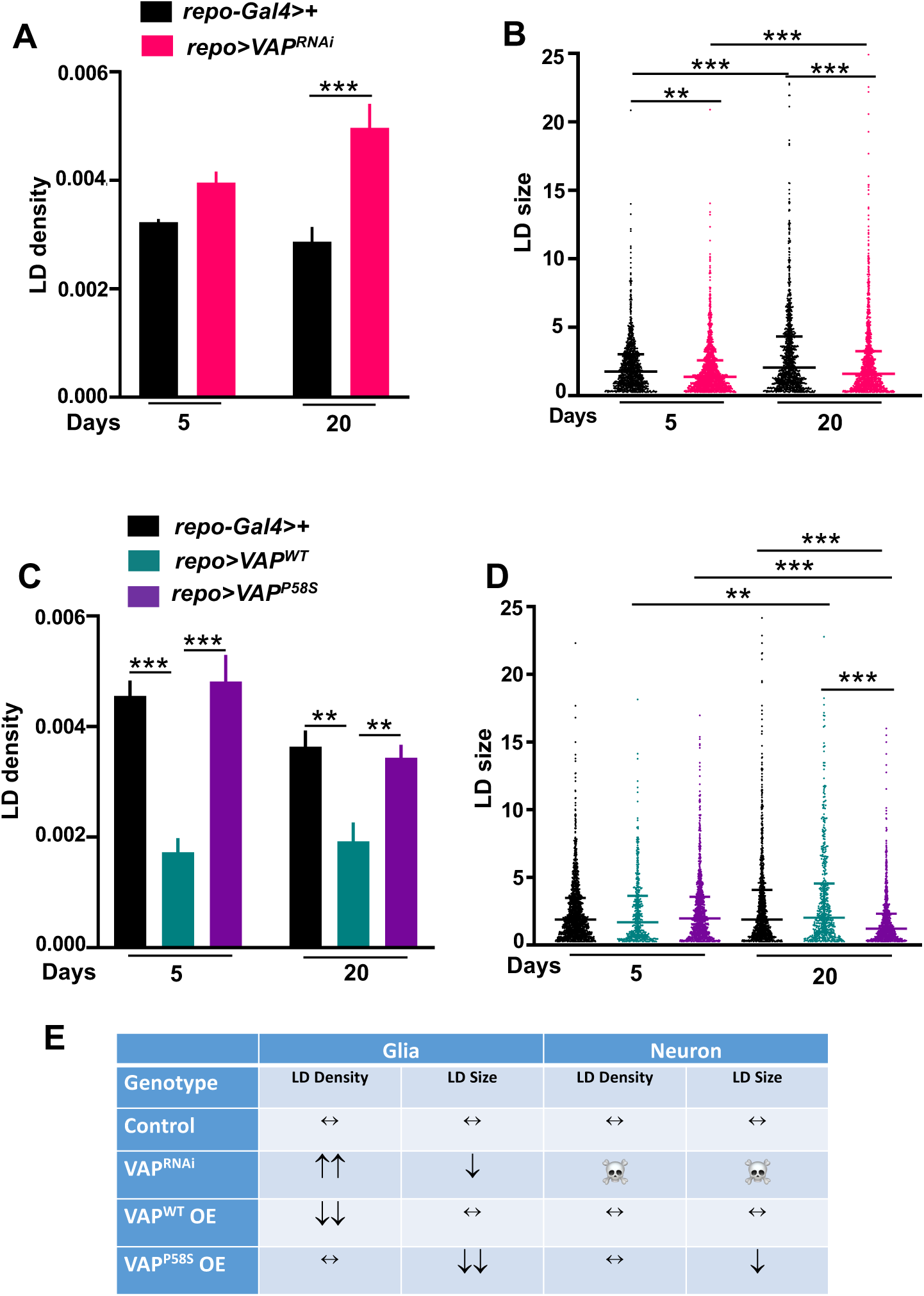
*VAP* modulation affects LD homeostasis in the *Drosophila* brain. **(A)** *VAP* knockdown in the glia leads to an age-dependent and significant increase in the LD density of the wild-type male brain tissue. Mean±SEM density is plotted on the Y axis in the graph against age (days) on the X axis. **(B)** Scattered dot plot is shown for LD size against age (days) on X axis for control and *VAP* knockdown in glia along with median and interquartile range value. N=5-6 adult brains, n=5 ROIs for each brain. **(C)** Overexpression of *VAP^WT^* in the glial cells of *Drosophila* cause significant decrease in LD density on 5^th^ day only which remains decreased on 20^th^ day as well whereas*VAP^P58S^* does not show any change. Quantitation of LD density (number of droplets per µm^3^) is plotted as mean±SEM in the graph. **(D)** Overexpression of *VAP^P58S^* in the glial cells of *Drosophila* leads to age-dependent decrease in LD size of the wild-type brain. Quantitation of LD size (volume in µm^3^) is plotted as scattered dot plot along with median and interquartile range. N=5 adult brains, n=5 ROIs for each brain. Statistical tests are performed for LD density using two-way ANOVA for mean comparison between genotypes of the same day, followed by Tukey’s test for multiple comparison correction, and Kruskal-Wallis test for LD size by comparing mean ranks between genotypes, followed by Dunn’s test for multiple comparison correction. p values are reported as *(<0.05), **(<0.01) and ***(<0.001). **(E)** Effect of VAP modulation in the glial and neuronal cells on the 20-day-old *Drosophila* brain LD is tabulated. VAP KD in neurons is developmentally lethal at 25 °C and this lethality can be overcome by reducing Gal4 activity by growing flies, till eclosion, at 18 °C (See Suppl. Fig. S6).

Next, we overexpressed (OE) *VAP^WT^* (*repo-Gal4>VAP^WT^*) and *VAP^P58S^* (*repo-Gal4> VAP^P58S^*) in glial cells and examined LD distribution of the adult brain for days 5 and 20, respectively (Fig. 5C-D; Suppl. Fig 5 C-E). Intriguingly, we found that upon glial *VAP^WT^* OE, the LD density was significantly reduced by over 60% on day 5 and remains low on day 20 (Fig. 5D). Interestingly, *VAP^P58S^*OE in glial cells did not affect LD density on either day. Median LD size remained unchanged for both *VAP^WT^* and *VAP^P58S^* when compared with the control on the 5^th^ day. However, *VAP^P58S^* OE showed a significant decrease (Fig. 5D) in the median LD size in 20-day-old flies, unlike control and *VAP^WT^* brains, where there was an increase in the size of LDs (Fig. 5D) with age (day-5 vs day-20).

We also tested *VAP KD* (*elav-Gal4>VAP^RNAi^*), OE of *VAP^WT^* (*elav-Gal4>VAP^WT^*) and *VAP^P58S^* (*elav-Gal4> VAP^P58S^*) in neurons and examined LD distribution in the brain. The neuronal data were compared with the glial data (Fig. 5E). VAP KD in neurons was lethal at 25 °C, suggesting that *VAP* activity in neurons is essential for development into adulthood. To gain a broad understanding of VAP function in neurons, we reduced VAP (*VAP KD*) during development by keeping *elav-Gal4>VAP^RNAi^* F1 embryos and larvae at 18 ℃ to minimize VAP KD (Chaplot *et al*. 2019) and then shifted flies to 25 ℃ post-eclosion. We saw a decrease in LD density and size on day 5, post eclosion (Suppl. Fig. 6 A-B), but these flies also started dying after day 10. For both neuronal *VAP^WT^* and *VAP^P58S^* OE (25 ℃ throughout), we observed a significant reduction in density on day-5, but by day-20 both *VAP^WT^* and *VAP^P58S^* OE show no change when compared to each other or controls. Neuronal *VAP^P58S^*OE, however, does lead to a significant decrease in LD size on day 20 (Suppl. Fig. 6 C-G).

In terms of differentiating VAP function in glia vs neurons, we also measured the effect of modulating VAP activity on the animal’s ability to climb as a function of age (SING Assay; Suppl. Fig. 7). In both glia (Suppl. Fig. 7A) and neurons (Suppl. Fig. 7B), climbing activity was affected with a drop in the climbing index after day-20, suggesting that VAP activity is important in both tissues.

In summary, glial VAP is important for lipid homeostasis, with LDs as a useful cellular readout, and the disruption of lipid homeostasis in the *VAP^P58S^* brain results from effects of both neurons and glia.

### Larval LDs are also modulated with VAP *gof*/*lof*

The modulation of LD density by *VAP* was unexpected, and to increase confidence on our observations, we repeated the experiment in the 3^rd^ instar male larval brain (Suppl. Fig. 8), targeting glia. We chose five ROIs at the tip of VNC for LD quantitation, as shown (Suppl. Fig. 8). We found that *VAP* KD in glial cells (*repo-Gal4>VAP^RNAi^*) resulted in higher LD density than in the control (Suppl. Fig. 8E). On the contrary, *VAP^WT^* OE showed a significant reduction in the LD density in the VNC of the larval brain. *VAP^P58S^*OE in the glial cells did not result in any change in LD density. When we examined the size of the LDs present in the ROIs, no change was detected upon modulating *VAP* in the glia of wild-type flies (Suppl. Fig. 8F). The larval LD data increased our confidence in the adult brain data, with both larval and adult brain data taken together suggesting that LD size in the adult was set developmentally, at least in the case of VAP modulation of LD in glia.

The next obvious question was to examine whether VAP could modulate LD density and size in vertebrates. From here on, the term ‘*VAPB*’ refers to the mammalian gene. We performed experiments on human-derived cell lines, repeating *VAPB KD*, *VAPB^WT^ OE,* and *VAPB^P56S^ OE* and examining changes in LDs.

### LDs in U2OS cells phenocopy the effects seen with VAP modulation in glia

Modulation of LDs by VAP could be *Drosophila-*specific and would be of greater significance if observed in vertebrates. To test this, we conducted equivalent experiments in the U2OS (human osteosarcoma) cell line (Fig. 6). For the *lof* experiment, we used a single interfering (si) RNA to KD *VAPB* in cells. The efficiency of KD is >95% (Fig. 6A), as confirmed by Western blotting. Next, we stained control and VAPB-KD cells with HCS LipidTOX™ Red Neutral Lipid Stain, imaged the cells (Fig. 6B) and then quantified the number and area of LDs per cell (Fig. 6C). As compared to the mock siControl, *KD* of *VAPB* led to a dramatic increase in both LD number and LD size (Fig. 6D). These results are in agreement with the data from the *Drosophila* brain, where *VAP KD* led to a shift in LD homeostasis, with higher LD number and increased LD size.

**Figure 6:**
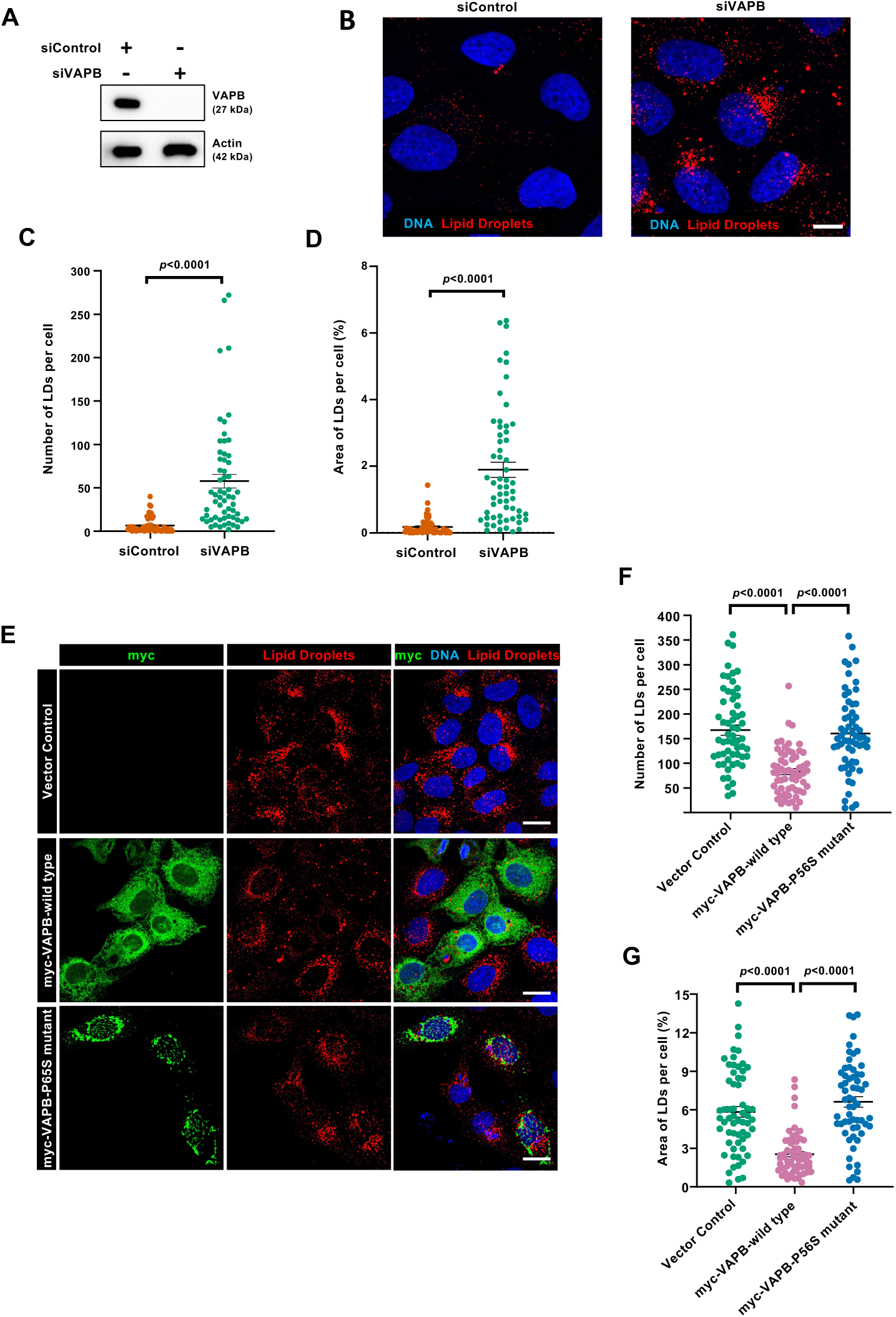
VAPB modulation leads to significant changes in LD size and density in U2OS cells. **(A-D)** U2OS cells were transfected with control (siControl) or VAPB-specific (siVAPB) siRNA for 72 h. (**A**) The extent of VAPB depletion by VAPB-specific siRNA was analysed by western blotting. (**B**) Confocal microscopic images of U2OS cells showing lipid droplets (LDs, red), detected by HCS LipidTOX™ Red Neutral Lipid Stain. DNA (blue) is visualized by Hoechst 33342 staining, Scale Bar, 20 μm. **(C)** The number of LDs per cell was quantified and plotted (*n* = 60 cells from three independent experiments). Data represented as mean±SEM, Student’s *t* test. *P* values are indicated. **(D)** The area of LDs per cell was quantified and expressed as percentage (%) of the cell area (*n* = 60 cells from three independent experiments). Data represented as mean±SEM, Student’s *t* test. *P* values are indicated. **(E-G)** Ectopic expression of VAPB^WT^, but not VAPB^P56S^ mutant, decreases LDs. U2OS cells were transfected with vector control, myc-VAPB wild-type or myc-VAPB P56S mutant for 56 h and supplemented with (50 μM) oleic acid (to induce LDs) for the last 8 h of transfection. **(E)** Cells were stained by myc-specific antibody to detect myc-tagged proteins (green), LDs by HCS LipidTOX™ Red Neutral Lipid Stain (red), and DNA was visualized by Hoechst 33342 (blue). Scale Bar, 20 μm. **(F)** The number of LDs per cell was quantified and plotted (*n* = 60 cells from three independent experiments). Data represented as mean±SEM, One way-ANOVA, Tukey’s multiple comparison test. *P* values are indicated. **(G)** The area of LDs per cell was quantified and expressed as percentage (%) of the cell area (*n* = 60 cells from three independent experiments). Data represented as mean±SEM, One way-ANOVA, Tukey’s multiple comparison test. *P* values are indicated.

Next, myc-tagged VAPB wild-type (myc-VAPB^WT^) or mutant (myc-VAPB^P56S^) was OE in U2OS cells (Fig. 6E). To induce LD formation, oleic acid was added to the cells in the last 8 h. In the case of VAPB^WT^ OE, the LD number per cell and LD size per cell were significantly decreased as compared to vector control expressing cells (Fig. 6 F, G). In contrast, *VAPB^P56S^* OE did not have any effect on the number or size of LDs (Fig. 6F-G) when compared to vector control.

Thus, from both fly and mammalian data, it is evident that cells containing LDs (Glia cells in *Drosophila* and U2OS cells treated with oleic acid) were susceptible to VAPB modulation. Reduction in VAP increased LDs, while OE of *VAP*/*VAPB* reduced LDs, suggesting that VAPB has a direct effect on LD dynamics. In contrast, OE of the mutant allele (*VAP^P58S^* or *VAPB^P56S^*), in the presence of *VAPB^WT^*, did not affect LDs. However, *VAP^P58S^* expression, in the absence of *VAP^WT^*, showed reduced LD density and an inability of LDs to grow into larger sizes with age.

## Discussion

Neurons and glia work together as metabolic partners to regulate normal brain function. This dynamic relationship includes two-way communication and the exchange of molecules (Bezzi and Volterra 2001; Fields and Stevens-Graham 2002; Allen and Lyons 2018).

This non-exhaustive list includes neurotransmitters such as glutamate, adenosine/ATP, GABA; energy currency such as lactate, glucose; Ions, including Na+, K+, Ca^2+^; growth factors, BDNF, NGF, TNF, interleukins; lipids such as fatty acids, cholesterol, phospholipids. In the specific case of lipids, glia act as a recycling center for neuronal lipids, receiving lipids exported by neurons and resupplying these in more useful forms. Glia also synthesize, from scratch, lipids based on neuronal demand. Thus, lipid homeostasis in the brain is centered around a healthy and dynamic neuro-glial axis (Ioannou *et al*. 2019; Islimye *et al*. 2022). Many recent reviews have highlighted roles for lipids in the brain and the consequences of lipid imbalance (Barber and Raben 2019; Yang *et al*. 2022; Byrns *et al*. 2024; Islam *et al*. 2025; Zhang *et al*. 2025).

Any deviation from the norm in the physiology of neurons or glia increases the load on their neighbouring, interconnected cells, e.g., neuronal stress can be relieved and balanced by glial action, but under chronic, long-term stress or disease conditions, the neuro-glial axis breaks down, leading to local or global malfunction or cell death (Farmer *et al*. 2020; Bohnert and Schrul 2024). Lipid homeostasis is a major metabolic process in the brain, given that the brain is a major lipid-storage organ, with LDs present in large amounts in brain tissue and glia as major lipid depots (Smolic *et al*. 2021; Teixeira *et al*. 2021; Islimye *et al*. 2022; Zhang *et al*. 2025). The majority of the lipids are synthesized at the ER membrane and then transferred to the other parts of the cell via membrane contact sites (Perry and Ridgway 2006; Mencarelli *et al*. 2010; Weber-boyvat *et al*. 2015; Cockcroft *et al*. 2016; Freyre *et al*. 2019; Xu *et al*. 2020; Kors *et al*. 2022a; Liu *et al*. 2023). Any perturbation in these sites affects ER lipid homeostasis and causes cellular dysfunctions (Perry and Ridgway 2006; Mencarelli and Martinez-Martinez 2013; Cockcroft *et al*. 2016; Gunay *et al*. 2021; Hartmann *et al*. 2022; Kim *et al*. 2023; Liu *et al*. 2023; Bezawork-geleta *et al*. 2025). VAP interacts with lipid transfer proteins to transfer lipids from the ER membrane to other parts of the cell, making it a crucial protein to study in the context of MCSs.

We are interested in lipid dys-equilibrium in the specific case of ALS. In the past few years, it has become increasingly clear that disturbances in lipid homeostasis are a signature of neurodegenerative disease, including ALS. Dyslipidemia, both hypo and hyper, is a feature of ALS patients (Phan *et al*. 2023; Cao *et al*. 2026). Commonly seen is the elevation of diglycerides, cholesterol, low-density lipoproteins and triglycerides, with lowered phospholipids, including PS and PC (Chelstowska *et al*. 2021; Phan *et al*. 2023). Some mutants of ALS loci such as *TDP43* and *FUS* show LD accumulation (Garcia-toledo *et al*. 2025; Long *et al*. 2026; Marcadet *et al*. 2026). Lower *VAPB* levels have been reported in other neurodegenerative models, which may contribute to LD accumulation (Girard *et al*. 2020). Inflammation is also a feature of lipid imbalance and of neuroinflammatory disease (Bernaus *et al*. 2020).

In our study, we focus on *ALS8* and measure dyslipidemia in the brains of flies carrying a *VAP^P58S^*mutation, which is analogous to pathogenic human VAPB mutation (*VAPB^P56S^)*. The mutant protein (VAP^P58S^ or VAPB^P56S^) aggregates (Kanekura *et al*. 2006; Teuling *et al*. 2007; Ratnaparkhi *et al*. 2008; Suzuki *et al*. 2009), modulates ER structure (Fasana *et al*. 2010; Papiani *et al*. 2012), causes ER stress (Kanekura *et al*. 2009; Moustaqim-barrette *et al*. 2014; Thulasidharan *et al*. 2024), and is haplosufficient in terms of development of the organism (Moustaqim-barrette *et al*. 2014), but insufficient in terms of maintaining cellular homeostasis as the organism ages (Moustaqim-barrette *et al*. 2014; Kulkarni *et al*. 2026). This suggests that *prima-facie*, VAP^P58S^ can function at par with VAP^WT^, with a significant amount able to fold and integrate in the ER membrane, while a large fraction is found in large, perinuclear (presumably inactive or non-functional) aggregates (Chaplot *et al*. 2019; Thulasidharan *et al*. 2024). Using mass spectrometry of brain lysates, we uncovered age-dependent lipid changes that mirrored the progression of motor dysfunction in the animal. In ageing flies, neutral lipids like CE and TAGs, increased with a concomitant decrease in cholesterol. Intriguingly, we did not detect significant (or large-scale) changes in phospholipid or glycolipid levels. In the adult brain, LDs were imaged and used to monitor LD density and size, which were again deviant from the norm. *VAP^P58S^* brains had lower LD density, which trended downward with age. Along the same lines, *VAP^P58S^* LD size was smaller and did not improve with age. Both cellular and physiological features of dyslipidemia in *VAP^P58S^*could be normalised by the addition of one copy of *VAP^WT^*. To relate lipid changes in *VAP^P58S^* animals with *VAP^WT^*function, we examined the relationship between VAP activity and its effect on lipid homeostasis. We found that excess VAP (and presumably excess VAP activity) in glia led to a dramatic decrease in LD density, without a change in LD size, while *VAP KD* led to an increase in LD density and a drop in LD size. *VAP^P58S^* OE did not significantly affect these trends. Similar results were seen by VAP modulation in human cells in culture. In comparison, roles for VAP in neurons appeared distinct from those in glia, with VAP *KD* leading to fewer, smaller LDs. In neurons, *VAP^P58S^* OE, unlike *VAP^WT^* OE led to smaller LDs. This suggests that the disruption of lipid homeostasis in the brain is probably an integrated function involving both neurons and glia, with brain LDs susceptible to VAP activity in both cell types (Fig. 7A).

**Figure 7:**
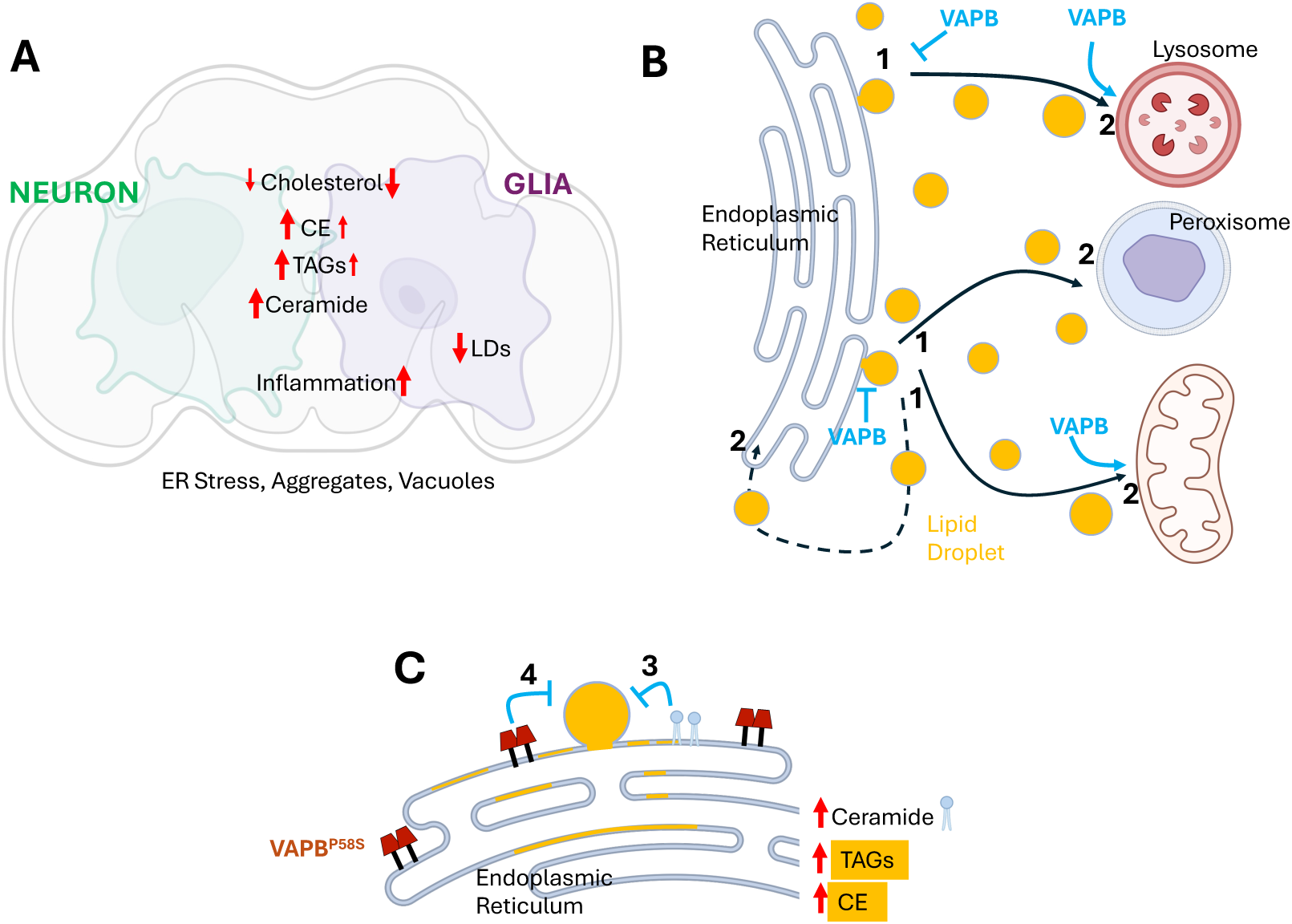
Age-dependent disruption of Lipid homeostasis is observed in a *Drosophila* model of ALS8. **(A)** Lipid Imbalance in *VAP^P58S^*appears to be a function of the neuro-glial axis, with an age-dependent reduction in LDs conflicting with increased CE & TAGs. We hypothesise that VAP plays critical and differential roles in both neurons and glia. LD and cholesterol drop are mostly glial, with increases in TAGs, CEs, and Ceramides mostly neuronal. Our results are consistent with earlier research implicating one cell type (glia) in major roles in inflammation (Kulkarni *et al*. 2026) and the other (neurons) in handling VAP^P58S^ aggregates (Moustaqim-barrette *et al*. 2014; Thulasidharan *et al*. 2024). **(B)** *VAPB^WT^* influences either LD formation or dissolution, or both. An increase in VAPB levels reduces LD biogenesis at the ER (1), or alternatively, increases LD utilisation (2) at the LD target/destination. **(C)** The Flux of LDs is modified in the *VAPB^P56S^* brain. Ceramides accumulate, modifying ER fluidity. This appears to affect the rate of LD production at the ER (3). Further, excess neutral lipids such as CEs and TAGs (marked in gold color) accumulate at the ER. *VAPB^P58S^* may also reduce the rate of LD production (4). Background schematics were generated using *Biorender*.

How does the *VAP^P58S^* allele disrupt lipid homeostasis, based on our understanding of VAPB’s roles in lipid metabolism? Since VAP decorates the cytoplasmic face of the ER and is also an integral part of membrane-membrane contact sites, its influence is, in all probability, either (a) based on its location and function as a tether and/or (b) its physical interaction with lipid transfer proteins. The simplest explanation (Model 1, Fig. 7B) would argue that, since LDs form at the ER, in the context of LD biogenesis, VAP may have a suppressive role; VAPs presence at the ER might decrease the rate of LD formation. Excess VAP might reduce LD biogenesis, whereas reduced VAP at the ER increases the rate of LD production. A second model (Model 2) would place VAP as a regulator of LD dissolution at destination compartments such as mitochondria, peroxisomes, or lysosomes. Here, VAP would function as an enhancer of LD clearance, with excess VAP leading to enhanced clearance, while reduced VAP would lead to excess LDs that pile up in the absence of clearance. This phenomenon has also been highlighted in a recent preprint, where *VAPB* loss leads to accumulation of CEs and TAGs along with LD accumulation (Borst Pauwels *et al*. 2026).

How would *VAP^P58S^*, the *ALS8* allele, then function in the cell in contrast to *VAPB^WT^*? *VAP^P58S^*may function as a gain-of-function allele, with LD numbers and size reduced in homozygous animals, in the absence of *VAP^WT^*. Based on Model 1, *VAP^P58S^* would then be a stronger suppressor of LD biogenesis. Previous studies, however, do not align with this definition. *VAP^P58S^* is a partially functional allele, is aggregation-prone, and can substitute for *VAP^WT^* function (Deivasigamani *et al*. 2014; Moustaqim-barrette *et al*. 2014; Thulasidharan *et al*. 2024).

Thus, we hypothesize (Fig. 7C) that the lack of correlation between excess TAGs & CEs at the ER, with the fewer LD’s in *VAP^P58S^* is *primarily* because of excess ceramides and low cholesterol affecting the fluidity and architecture of the ER membrane.

The excess ceramides reduce the ER membrane’s ability to produce LDs efficiently (Fig. 7C). The clustered VAP^P58S^ on the ER membrane may also influence core proteins that assist in LD biogenesis (Fig. 7C). VAP^P58S^ also causes ER stress (Kanekura *et al*. 2009; Suzuki *et al*. 2009; Thulasidharan *et al*. 2024), which could also be a contributory factor. These features would also explain the inability of LDs to grow in size in the *VAP^P58S^* mutant, as shown in an earlier study of preadipocytes (Tokutake *et al*. 2015).

The exact mechanism through which VAP regulates lipid balance in the cell needs to be elucidated. The search for the mechanism is complicated by VAP’s ability to interact with >350 human protein (Huttlin *et al*. 2015; Olkkonen 2015), a large proportion of which are involved in lipid trafficking (Perry and Ridgway 2006)(Murphy and Levine 2016). VAP’s influence may be through one or many of these interactors. For example, we have recently demonstrated that Ceramide Transfer Protein (CERT), a VAP interactor, can influence lipid balance and LD formation (Garg *et al*. 2026). Further, as a tethering protein, VAP can affect the architecture of the MCS (Kors *et al*. 2022a; Kalarikkal *et al*. 2024; Kors *et al*. 2024; Obara *et al*. 2024).

MCS changes between organelles would also influence lipid flux, adding another layer to lipid regulation (Quon and Beh 2015).

In summary, our study, using both *Drosophila* and mammalian disease models, uncovers an imbalance of lipid metabolism in *ALS8*. The disruption appears to be due to impairment in the neuron-glia axis, with LDs in glia serving as a cellular marker of disease. LDs are sensitive to VAP/VAPB levels in the glia, highlighting their roles as important metabolic partners for neurons. LD flux also depends on VAP/VAPB function, with our data suggesting that VAP regulates LD size and density by modulating LD biogenesis at the ER and/or LD uptake or dissolution.

## Materials & Methods

### *Drosophila* husbandry, fly stocks and reagents

All experimental stocks were maintained on standard cornmeal agar medium, and crosses were set up in a 12 hr Light: 12 hr Dark cycle at 25℃ unless specified. *Canton-S* is used as wild-type control in the experiments. *ΔVAP(Δ166)*, *gVAP^P58S^*, *gVAP^WT^* were kind gifts from the Tsuda lab and are described earlier (Moustaqim-barrette *et al*. 2014; Tendulkar *et al*. 2022; Thulasidharan *et al*. 2024). *ΔVAP;;gVAP^P58S^* was used as an *ALS8* model, *ΔVAP;;gVAP^WT^*/*Tb*, *ΔVAP;repo-Gal4;gVAP^P58S^* are described in (Tendulkar *et al*. 2022; Thulasidharan *et al*. 2024). *repo-Gal4* was a kind gift from Dr. Bradley Jones. *elav-Gal4* and has been described in (Ratnaparkhi *et al*. 2008). Bloomington Stocks: *Canton-S* (BL_0001), *VAP^RNAi^* (BL_77440), were used for experiments. *UAS-VAP^WT^* and *UAS-VAP^P58S^*transgene lines are described in (Ratnaparkhi *et al*. 2008; Chaplot *et al*. 2019).

### Lipid extraction and targeted LC-MS lipidomics

For sample preparation, *Drosophila* adult brains were dissected in ice cold 1X PBS with 2 brains in each vial and stored at -80℃ after flash freeze with liquid N_2_. Then, lipids were extracted with MS grade organic solvents using the Folch extraction method as described (Pathak *et al*. 2018). Internal standards from Avanti polar lipids were used as follows: 1nmol each of Cholesterol d7, 17:1 Sphingosine, 17:1 free fatty acid, 37:4 Phophatidylcholine, 37:4 Phosphatidylethanolamine, 37:4 Phosphatidic acid, 37:4 Phosphatidylglycerol, 17:0/34:1 Triacylglycerols, 32:0 d5-glycerol and 100 pmol each of 17:1 Sphingosine-1-phosphate, 25:0 Ceramide, 12:0 Sphingomyelin, 20:4 d5-glycerol, 19:0 Cholesterol ester, 12:0 Ceramide-1-phosphate, 37:4 phosphatidylserine, phosphatidylinositol, 17:1 Lyso-PC, 17:1 Lyso-PE, 17:1 Lyso-PA, 17:1 Lyso-PG, 17:1 Lyso-PS, 17:1 Lyso-PI. After the successful extraction, lipids were resolubilized in 200 ul of 2:1(v/v) of CHCl_3_, Me-OH and 20 µl was used for injection. All the lipid species were quantified as per protocol described (Pathak *et al*. 2018). Phospholipid species were quantified using multiple reaction monitoring high resolution (MRM-HR) scanning method on a Sciex X500R QTOF LC-MS with an Exion-LC series quaternary pump. All the data was acquired and processed through Sciex-OS software.

### Lipid Droplet staining for larval and adult brains

Late 3^rd^ instar wandering male larvae were selected for the study where brains were dissected in ice cold 1X PBS. Fixation is done for 20 minutes with 4% PFA (0.3% Triton-x-100) followed by two immediate washes of 1X-PBS and then 3^rd^ wash of 15 minutes with 1 X PBS at room temperature. 1:1000 BODIPY^TM^ 493/503 (1mg/ml, Invitrogen, D3922) along with 1:1000 Hoechst (10mg/ml, Invitrogen) in 1X PBS were added to the brains and incubated on rocker for 30 minutes at room temperature under dark conditions. Then, solution was removed and brains were rinsed with 1X PBS. Immediate mounting was done for 5-6 brain samples using slowfade mounting media (Vectashield S36937). For adult brains, protocol was adapted from (Garg *et al*. 2026) and proceeded further with imaging and analysis.

### Microscopy imaging and analysis

Mounted samples (both larval and adult brains) were imaged on Leica SP8 with 63X objective lens as described (Garg *et al*. 2026). Laser power and power gain were adjusted based on individual experiments but kept constant across genotypes and age. Fiji software was used for LD quantification. Firstly, mean intensity value of each slice was subtracted from that slice of the stack using a macro code. For larval brains, five ROIs of equal area were taken for analysis. For adult brains, Five Regions of Interest (ROIs) were drawn near the cells surrounding SOG region for analyzing LDs. Quantitative analysis for LD density and LD size was performed using macro code as described (Garg *et al*. 2026). For LD density, total number of droplets in each ROI were divided by volume of ROI based on nuclei stack resulting number of droplets per unit volume of µm^3^. LD size (volume in µm^3^) is plotted in the form of scattered dot plot by taking size of all droplets in the ROIs of total brains of each genotype into consideration (Garg *et al*. 2026).

### Climbing Assay

Climbing activity of each genotype was observed using the SING assay as described (Garg *et al*. 2026). Briefly, three biological replicates of each genotype with 30 age-matched male flies were transferred to a 250 ml glass cylinder and allowed to acclimatize for 5 minutes. Then, cylinder was tapped for three times to startle the flies and three readings were taken after 1 minute for each genotype with 3 minutes of resting period after each test. After 1 minute, flies were counted based on three classes, non-climbers, poor climbers (0-80ml, 7.5cm) and good climbers (above 80ml or 7.5cm). These classes were given score of 0, 1 and 2. The experiment was done after every five days from eclosion until it was not required. CO_2_ was used to anaesthetise the flies after 3 trials were done, and they were transferred to fresh media vials. Climbing Index was calculated by multiplying each score by the number of flies in that range, and the sum of all three scores was divided by the total number of flies for each genotype. Two-way ANOVA was used as a statistical test, followed by Tukey’s test for multiple comparisons between genotypes.

### Mammalian Cell Culture, siRNA and transfections

U2OS cells were used for all experiments and were maintained in DMEM (10569-010, Gibco) containing 10% FBS (F2442-500ML, Sigma Aldrich) supplemented with 10 μg/ml ciprofloxacin (Cipla). The cells were routinely checked for mycoplasma contamination. pCI-neo-myc-VAPB WT and pCI-neo-myc-VAPB P56S DNA constructs were generous gifts from Christopher Miller (King’s College London, UK). The DNA constructs were transfected for 48 h using polyethyleneimine, linear (PEI, MW-25000; Polysciences Inc.) for the VAPB overexpression studies.

For VAPB depletion studies, the siRNAs were designed against the following target sequences: siControl (5’-TTCTCCGAACGTGTCACGT-3’)(Sahoo *et al*. 2017), siVAPB (5’-GCTTTCCGTGTCTTCAGTT -3’). Control siRNA was obtained from Dharmacon and VAPB-specific siRNA was procured from Sigma Aldrich (Cat. No. SASI_HS01_0019018).

Lipofectamine RNAimax (13778150, Invitrogen) was used for siRNA transfections, as per manufacturer’s protocol. The siRNAs were transfected for 72 h.

### U2OS Cell Lysis and Western Blotting

Cells were lysed in RIPA buffer [50 mM Tris-HCl (pH 7.4), 150 mM NaCl, 1% NP-40, 0.1% SDS, 0.5% sodium deoxycholate], supplemented with 50 mM NaF, 5 mM sodium pyrophosphate, 1 mM PMSF, 0.75 mM sodium orthovanadate, and protease inhibitor cocktail (Cat. No. 11836170001, Roche). After retro-pipetting thoroughly, cells were left on ice for 5 min, then centrifuged at 12,900 × *g* for 20 min to pellet cell debris, and the clear supernatant was collected. Protein concentration was estimated with the Bradford (Bio-Rad, Cat. No. 5000006) assay. Samples were separated using SDS-PAGE and transferred onto a PVDF membrane (Millipore) for western analysis. The PVDF membranes were probed with anti-VAPB antibodies (66191-1-Ig; Proteintech) at 1:3000 dilution, and anti-β-Actin antibodies (sc-47778; Santa Cruz Biotechnology) at 1:4000 dilution.

### U2OS Cell immunofluorescence (IF) imaging and Analysis

Cells seeded on coverslips were fixed by 4% PFA for 20 min, followed by washing once with phosphate-buffered saline (PBS, pH 7.4). Later, cells were permeabilized using 0.1% Triton-X-100 (Sigma) for 7 min. Then, cells were washed with PBS and incubated with mouse anti-c-myc antibody (sc-40) from Santa Cruz Biotechnology (1:500 dilution) in PBS containing 2% normal horse serum (NHS, Cat. No. S-2000-20, Vector Laboratories) for 45 min. After washing with PBS three times, cells were incubated with PBS containing 2% NHS with donkey anti-mouse Alexa Fluor 488 secondary antibody (A21202, Invitrogen, Thermo-Fisher) at 1:1000 dilution, HCS LipidTOX™ Red Neutral Lipid Stain dye (Cat. No.H34476, Invitrogen, Thermo-Fisher) at 1:1000 dilution, and Hoechst 33342 dye (Cat. No. B2261, Sigma). After three PBS washes, coverslips were mounted on glass slides with ProLong™ Glass Antifade Mountant (Cat. No. P36980, Invitrogen, Thermo-Fisher). Images were acquired using an Olympus FV3000 confocal microscope with a 60x oil objective (1.42 NA). LUTs were adjusted for the λ564 (red) channel with Min. 1800 and Max. 4095. The images were processed using cellSens software (Olympus) and converted into TIFF files for further analysis.

The nucleus and transmitted light (TD) TIFF images were imported into FIJI version 1.53 (ImageJ, NIH, US), merged and used to mark individual cell boundaries, recorded as independent regions of interest (ROI). These ROIs were then interpolated onto the Lipid droplet image [λ564 (red) channel] for quantifying LDs per cell. The pipeline order that was followed for quantitation is: Convert into 8-bit from RGB color > Convert to binary > Apply water shedding function (to segment clustered LDs) > Select the interpolated ROIs one by one > Use ‘analyze particles’ option with the following dimensions: Size (in μm^2^): 0.1 - infinity; Circularity: 0.00 – 10.00 and showed as overlaid masks with include holes options ticked in. This pipeline was repeated for all the ROIs selected for each image. The summary was exported as an Excel sheet with the following parameters included: LD count, total LD area, average size of LD and percentage area of LD in the cell. The parameters LD number and the percentage area of LD per cell were used for statistical analysis.

### Statistical tests

All statistical analyses were performed using either GraphPad Prism 8.0 or GraphPad 10 and were calculated from independently repeated experiments as indicated in the respective figure legends. Data represented as mean±SEM. For mammalian experiments, unpaired Student’s *t-*test was applied for the VAPB depletion study, and One way-ANOVA, Tukey’s multiple comparison test was applied for the VAPB overexpression study. *P* values were as indicated in the figures; *P* ≤ 0.05 were considered significant and *P* > 0.5 were not significant.

## Data & Resource availability

Raw imaging data will be shared on acceptance of the manuscript, as per journal policy.

## Contributions

GSR, LG, KC, JJ and SK conceptualized the project. LG, KC, and ST executed the *Drosophila* experiments, with GK contributing to the mammalian experiments. LG & GSR wrote the initial draft, and all authors contributed to the final draft. GSR, SK and JJ were responsible for funding acquisition, supervision and project management.

## Acknowledgements

We thank: Bloomington *Drosophila* Stock Centre (BDSC), supported by NIH grant P40OD018537, for fly stocks; Prof. Hugo Bellen & Dr. Bhagyashree Kaduskar for their feedback; Snehal Patil and Yashwant Pawar for fly media and stock maintenance; The microscopy facility at IISER Pune, managed by Dr Santosh Podder and Vijay Vitthal and at NCCS Pune managed by Dr. Ashwini Atre and Ms. Trupti Kulkarni, are gratefully acknowledged.

---

**Suppl. Figure 1.**
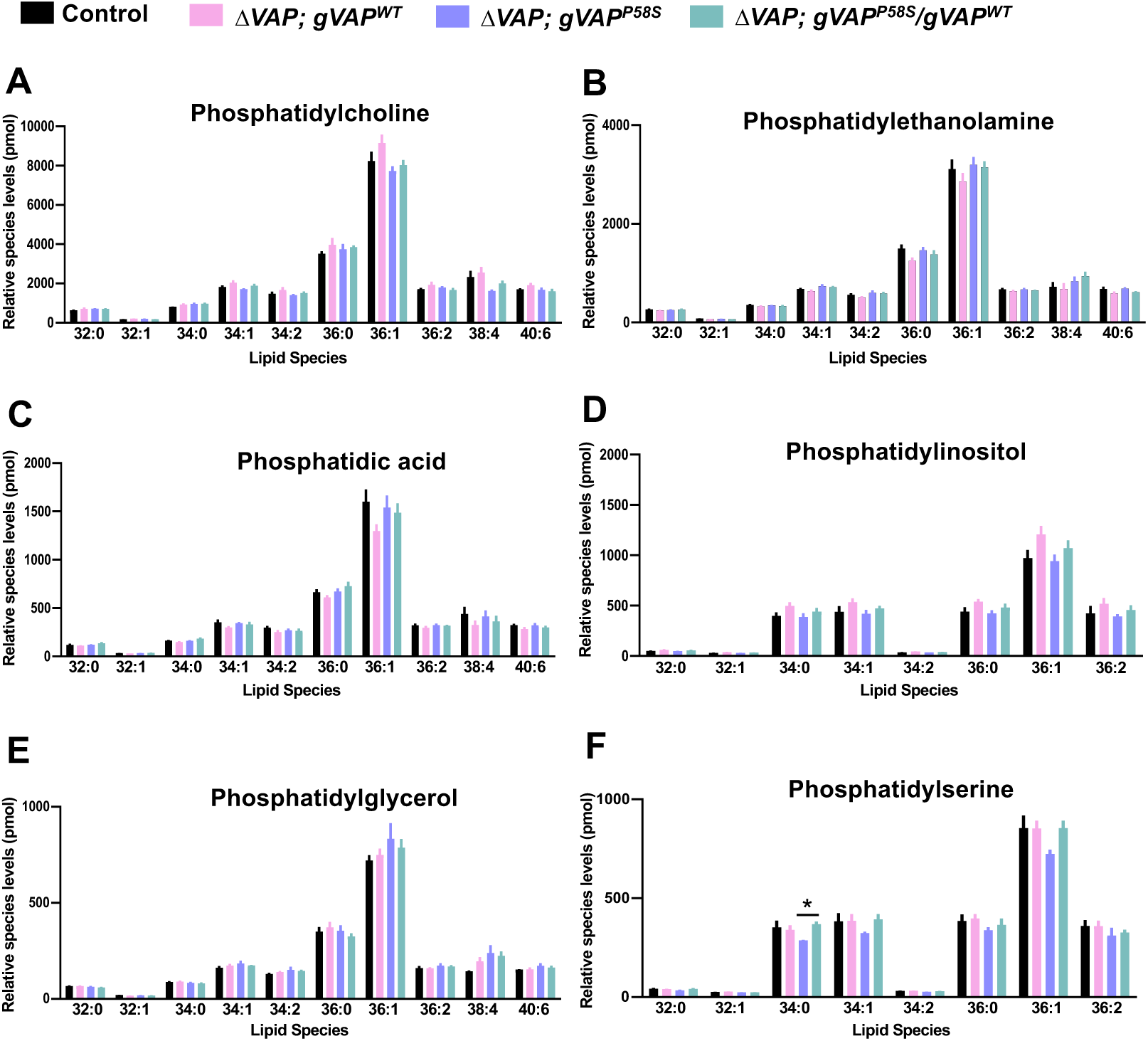
LC-MS lipid quantitation for phospholipids in 15 day old *Drosophila* male brain. No significant change is detected in the phospholipid species of *ΔVAP; gVAP^P58S^* compared to controls for (A) Phosphatidylcholine (PC), (B) Phosphatidylethanolamine (PE), (C) Phosphatidic acid (PA), (D) Phosphatidylinositol (PI), (E) Phosphatidylglycerol (PG). However, Phosphatidylserine(PS) C34:0 is significantly reduced in *ALS8* model which are rescued by single copy of *VAP* (F). Multiple unpaired t-test is performed for mean comparison between genotypes with FDR approach of Benjamin and Hochberg for multiple comparison correction with p values reported as *(<0.05), **(<0.01) and ***(<0.001). N=5 biological replicates, n=2 adult brains for each genotype.

**Suppl. Figure 2.**
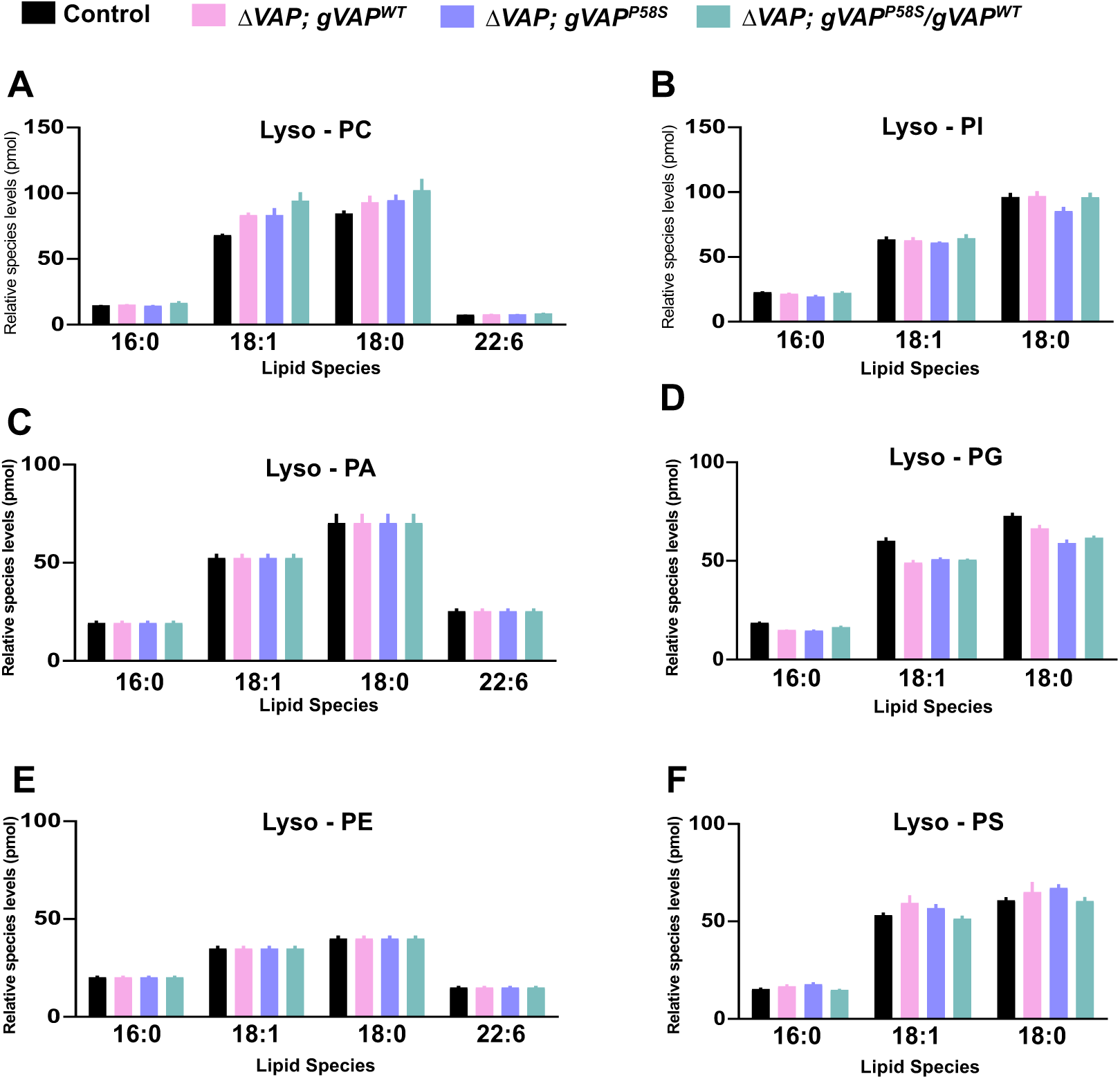
LC-MS lipid quantitation for lyso-phospholipids in the 15 day old *Drosophila* male brain. No significant change is detected in lyso-phospholipid levels of *ΔVAP; gVAP^P58S^* compared to controls for (A) Lyso-Phosphatidylcholine (Lyso-PC), (B) Lyso-Phophatidylinositol (Lyso-PI), (C) Lyso-Phosphatidic acid (Lyso-PA), (D) Lyso-Phosphatidylglycerol (Lyso-PG), (E) Lyso-Phosphatidylethanolamine (Lyso-PE) and (F) Lyso-Phosphatidylserine (Lyso-PS). Multiple unpaired t-test is used as statistical test for mean comparison between genotypes with FDR approach of Benjamin and Hochberg for multiple comparison correction with p values reported as *(<0.05), **(<0.01) and ***(<0.001). N=5 biological replicates, n=2 adult brains for each genotype.

**Suppl. Figure 3.**
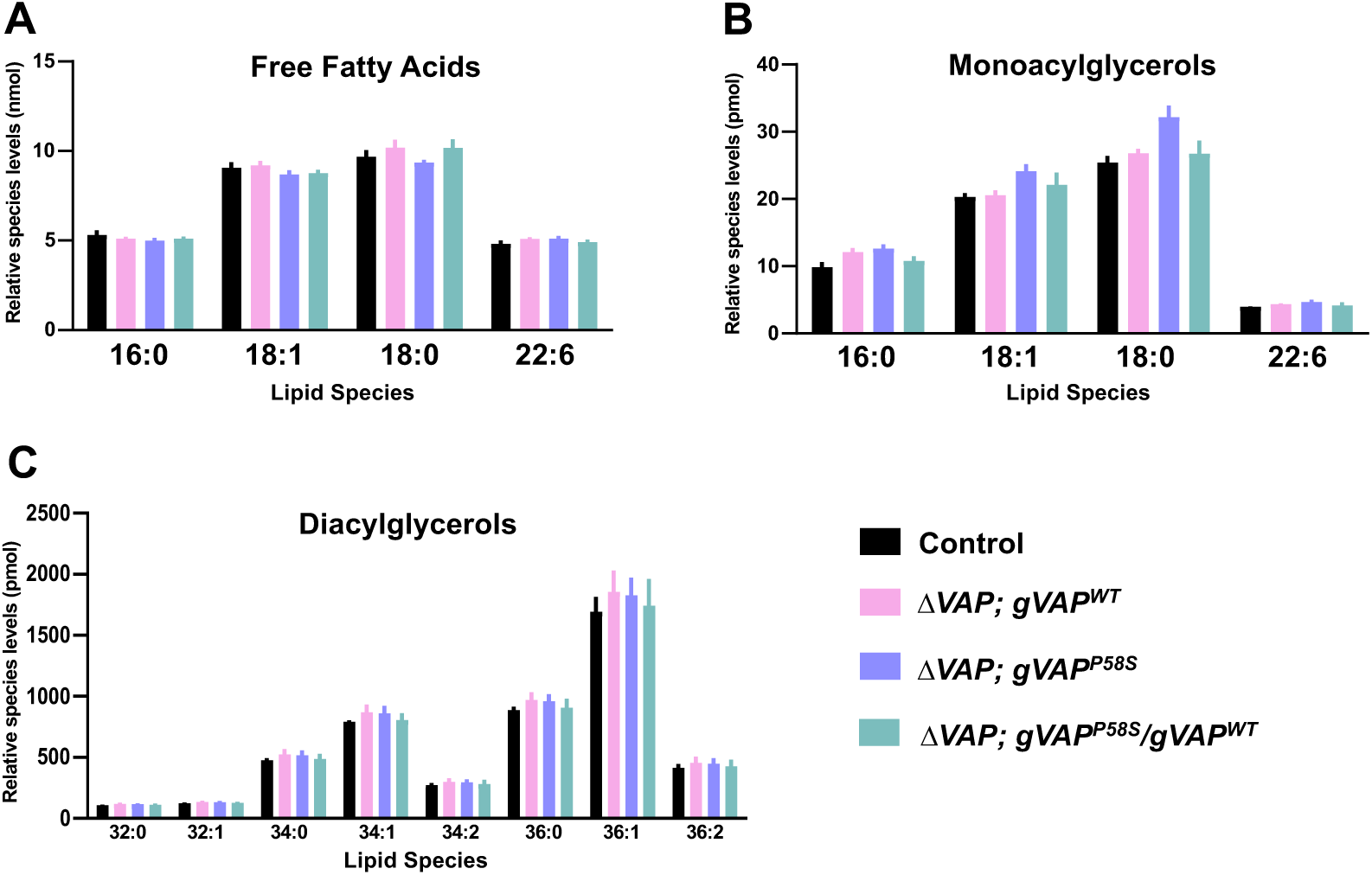
LC-MS lipid quantitation in the 15 day old *Drosophila* male brain. Relative lipid/species are plotted in the graph as mean±SEM for (A) Free Fatty Acids (FFA), (B) Monoacylglycerols (MAGs) and (C) Diacylglycerols (DAGs). Lipid levels remain unchanged for the mentioned speices in the brain of *ΔVAP; gVAP^P58S^* male flies at day 15 compared to controls. Multiple unpaired t-test is used as statistical test for mean comparison between genotypes with FDR approach of Benjamin and Hochberg for multiple comparison correction. N=5 biological replicates, n=2 adult brains for each genotype.

**Suppl. Figure 4.**
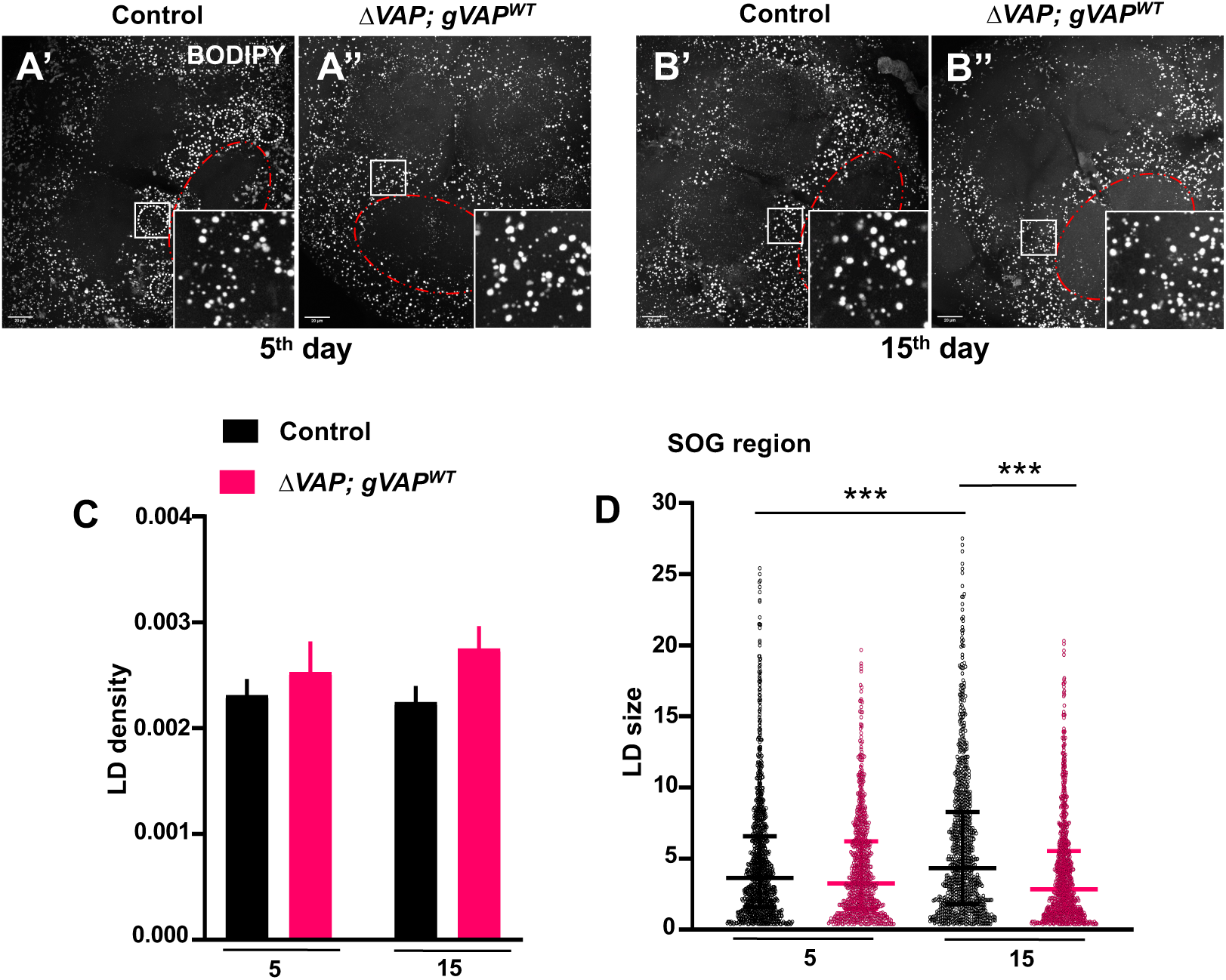
No change is detected in LD density of *ΔVAP; gVAP^WT^ Drosophila* adult brains. **(A-B)** Representative MIPs of BODIPY-stained LDs in the male adult brain’s central body with a zoomed inset shown in the lower right corner. Scale bar is shown as 20 µm. (A) LD distribution is shown at 5^th^ day for wild-type control (A’) and *ΔVAP; gVAP^WT^*(A’’). Similarly, (B) represents the distribution for 15-day-old male flies. SOG region is selected for the analysis of LDs depicted by red outline. ROIs are represented around SOG region in control (A) used for LD quantitation. **(C)** Quantitation for LD density (number of droplets per µm^3^) plotted as mean±SEM values on Y axis against age (days) on X axis. Two way ANOVA was performed for statistical test between genotypes followed by Tukey’s test for multiple comparison correction with p values reported as *(<0.05), **(<0.01) and ***(<0.001). **(D)** Scattered dot plot for LD size (volume, µm^3^) is plotted with size on the Y axis, including median with interquartile range, against age (days) on the X axis. Kruskal-Wallis test was used for mean rank comparison between genotypes, followed by Dunn’s test for multiple comparison correction with p values reported as *(<0.05), **(<0.01) and ***(<0.001). N=5 adult brains per genotype, n= 5 ROIs analyzed for each brain.

**Suppl. Figure 5.**
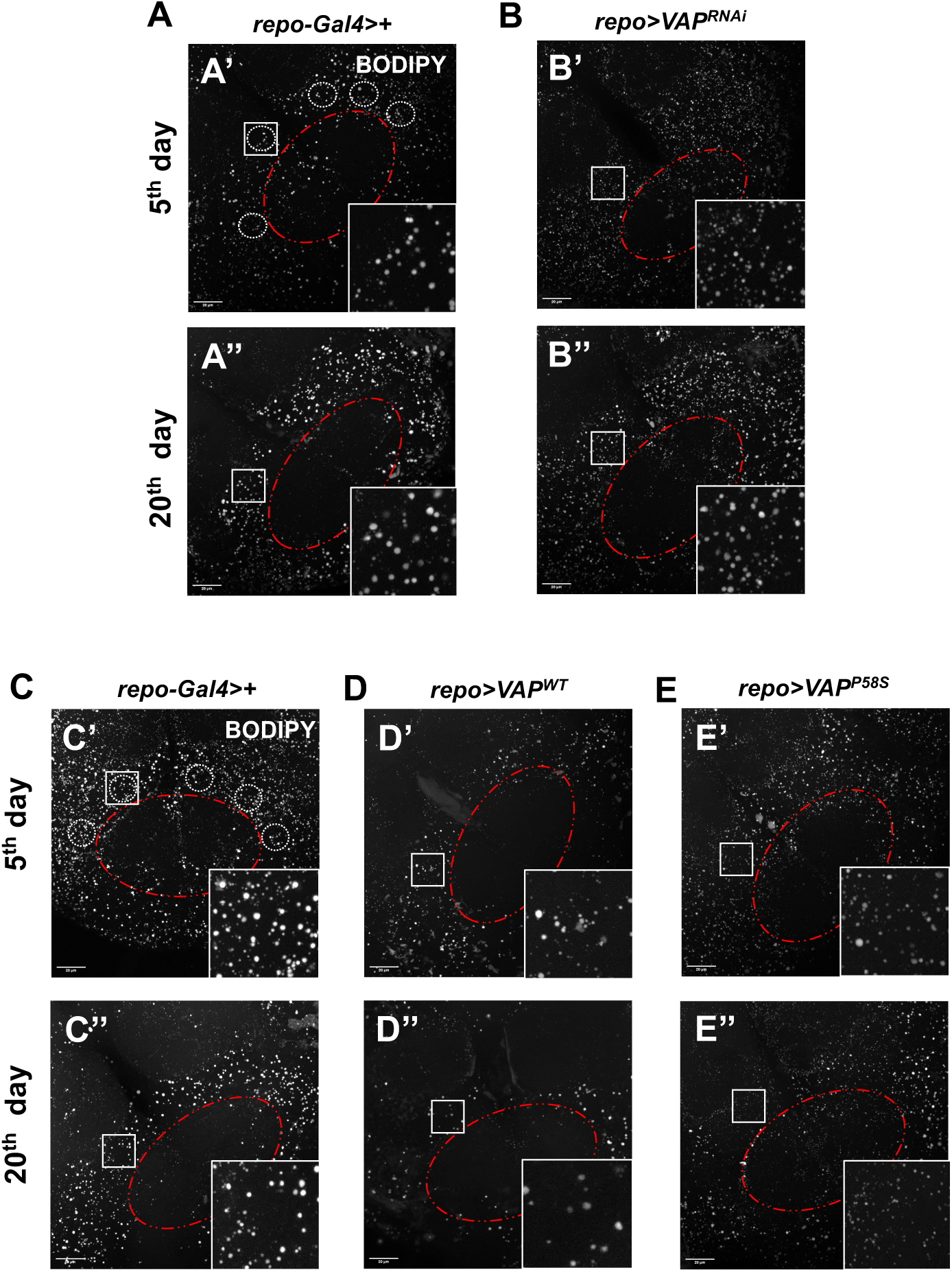
*VAP* is a negative regulator for LD distribution in the glial cells of *Drosophila*. **(A-B)** shows MIPs (20% Brightness) of BODIPY-stained adult brain SOG region, with zoomed inset on the lower right corner. Scale bar = 20µm. The SOG region is selected for LD analysis, as depicted by a red outline. ROIs are represented around the SOG region in the control (A’) used for LD quantitation. N=5-6 adult brains, n=5 ROIs for each brain. **(C-E)** are representative images (MIP, 20% Brightness) of BODIPY-stained LDs in the adult *Drosophila* brain (SOG region) of 5 and 20-day-old male flies following glial *VAP* modulation. Zoomed insets are shown in the lower right corner of each image. Scale bar is represented as 20µm. Five ROIs are drawn around SOG region (outlined by red color) in control (C’) used for LD analysis. N=5 adult brains, n=5 ROIs for each brain.

**Suppl Figure 6.**
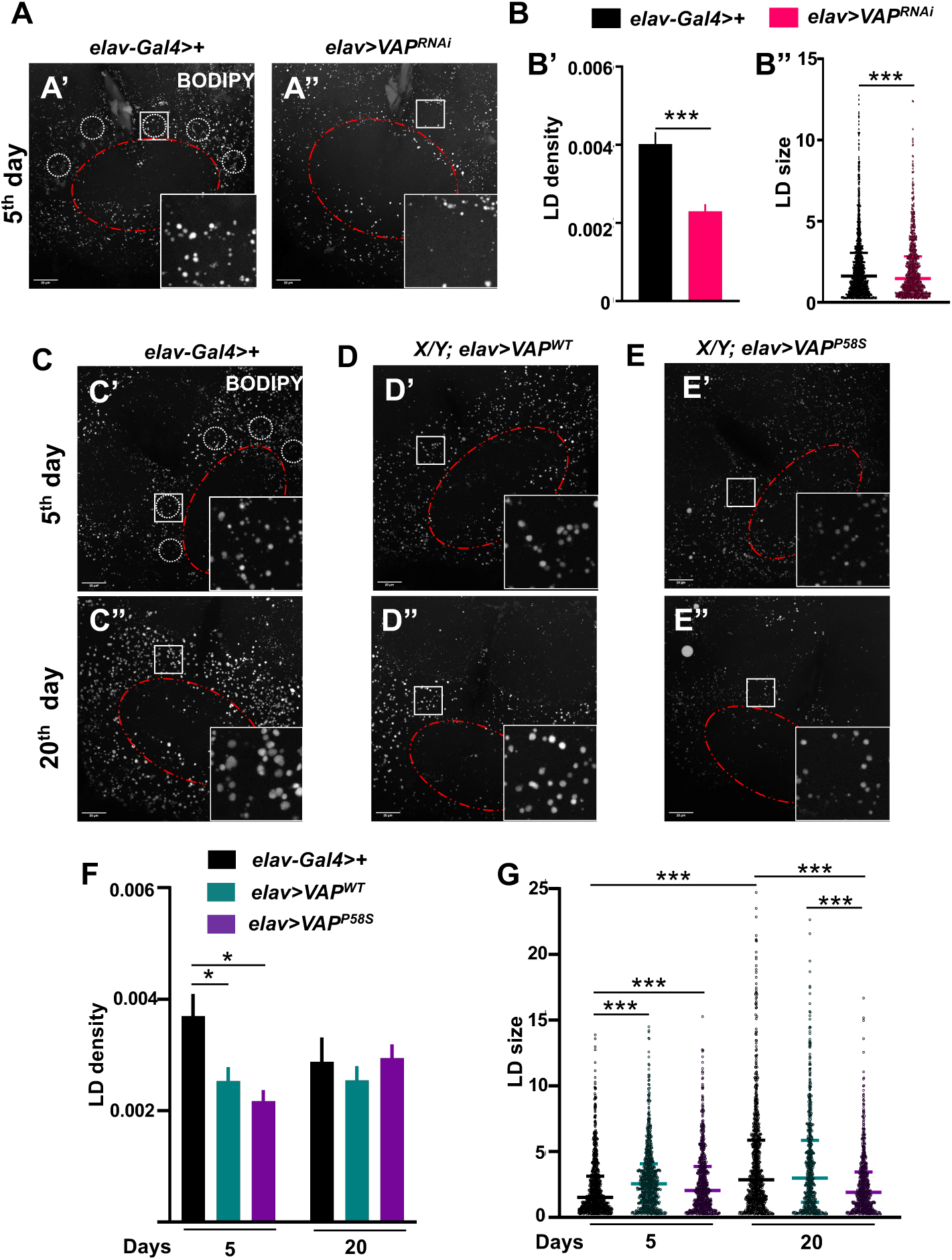
Effect of neuronal *VAP* modulation on the adult brain LD homeostasis. **(A)** shows MIPs (20% Brightness) of BODIPY stained LDs in the SOG region of the 5 day old *Drosophila* adult brain with zoomed inset on the lower right corner. Scale bar = 20µm. (A’) shows LD distribution in the adult brain for *elav-Gal4* control whereas (A’) shows reduced LD distribution on *VAP* KD in the neuronal cells of the male flies. Five ROIs are drawn around SOG region (outlined by red color) in control (A’) used for LD analysis. **(B)** Quantitation of LD distribution is plotted using LD density (number of LDs per µm^3^) and LD size (volume in µm^3^) as parameters. (B’) Mean±SEM value is plotted on Y axis as LD density. Unpaired t-test is used as statistical test for mean comparison between genotypes followed by Welch’s test with p values reported as *(<0.05), **(<0.01) and ***(<0.001). (B”) Scattered dot plot is shown for LD size for control and *VAP* knockdown in neurons along with median and interquartile range value. Non-parametric t-test for LD size is used for mean rank comparison followed by Mann Whitney test, p-values are mentioned using APA style. N=5-6 adult brains, n=5 ROIs for each brain. **(C-E)** are representative images (MIP, 20% Brightness) of BODIPY stained LDs in the adult *Drosophila* brain (SOG region) of 5 and 20 day males flies following neuronal *VAP* modulation. Zoomed insets are shown in the lower right corner of each image. Scale bar is represented as 20µm. Five ROIs are drawn around SOG region (outlined by red color) in control (C’) used for LD analysis. (C) shows images for Gal4 control of 5 day old (C’) and 20 day old (C”) brain samples. **(D)** represents images for LD distribution on overexpressing *VAP^WT^* in the neurons for 5^th^ day (D’) and 20^th^ day respt. (D”). **(E)** *VAP^P58S^* OE in the neurons of *Drosophila* leads to changes in LD size on day 5 (E’) as well as day 20 (E”) **(F)** Quantitation of LD density (number of droplets per µm^3^) is plotted as mean±SEM value in the graph. Two way ANOVA is used for mean comparison between genotypes of same day followed by Tukey’s test for multiple comparison correction with p values reported as *(<0.05), **(<0.01) and ***(<0.001). (G) Quantitation of LD size (volume in µm^3^) is plotted as scattered dot plot along with median and interquartile range. Kruskal Wallis test is used for mean rank comparison between genotypes followed by Dunn’s test for multiple comparison correction with p values reported as *(<0.05), **(<0.01) and ***(<0.001). N=5-6 adult brains, n=5 ROIs for each brain.

**Suppl. Figure 7.**
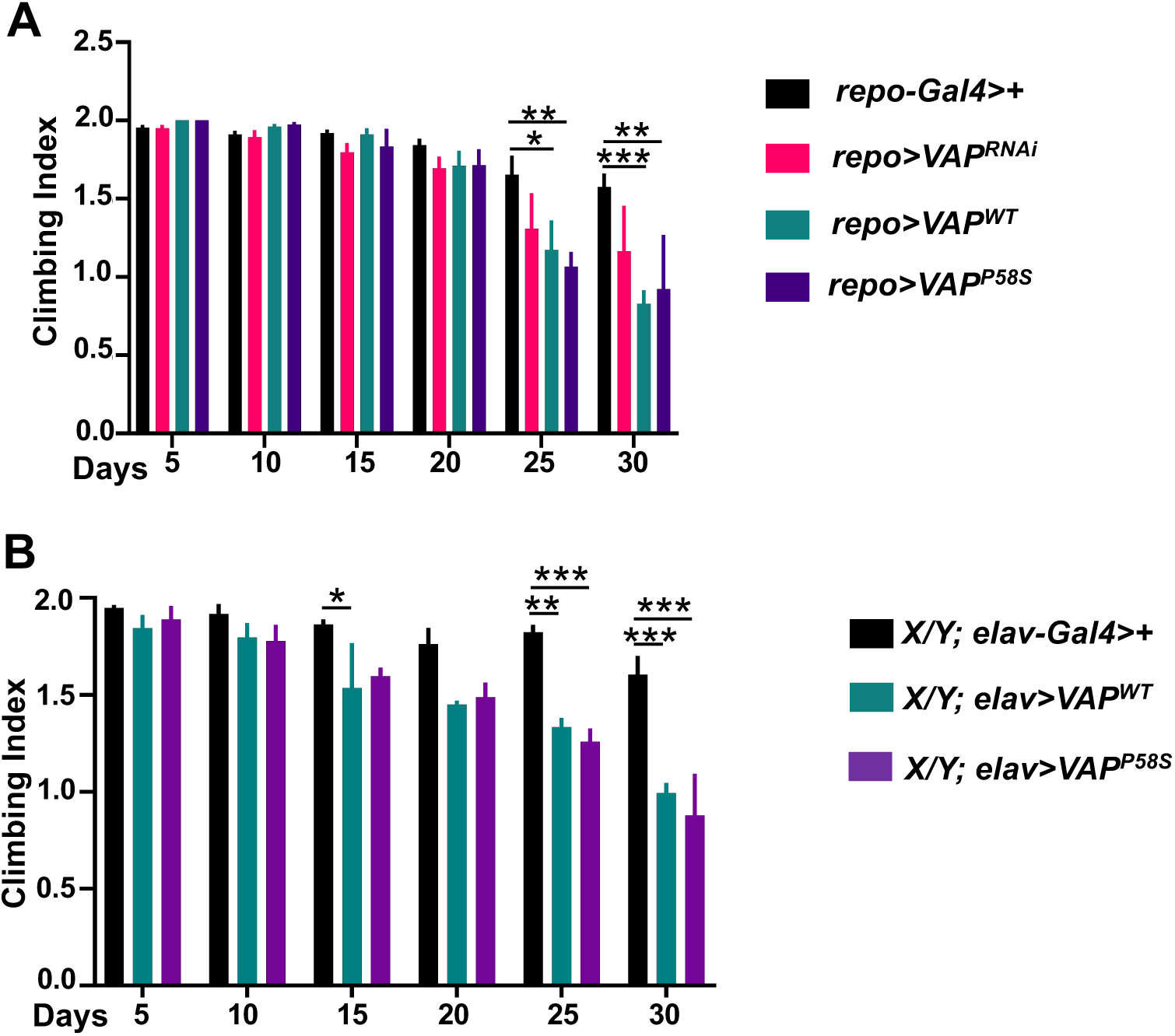
Effect of *VAP* modulation on the motor function of the flies. **(A)** Climbing Index is plotted for the motor function on modulating *VAP* in the glia of *Drosophila.* Removal of *VAP* in the glial cells does not appear to affect climbing activity of the flies, but more VAP, either wildtype or mutant (VAP^P58S^), leads to significant motor impairment day-25 onwards. **(B)** Climbing Index is plotted for the motor function on modulating *VAP* in the neurons of *Drosophila. VAP^WT^* OE in the neuronal cells leads to motor defects on day 15 onwards whereas *VAP^P58S^*OE leads to significant motor dysfunction on 25^th^ day onwards. Two-way ANOVA was performed for mean comparisons between genotypes on the same day, followed by Tukey’s test for multiple comparison correction with p values reported as *(<0.05), **(<0.01) and ***(<0.001). N=3 biological replicates, n∼30 male flies for each genotype.

**Suppl. Figure 8.**
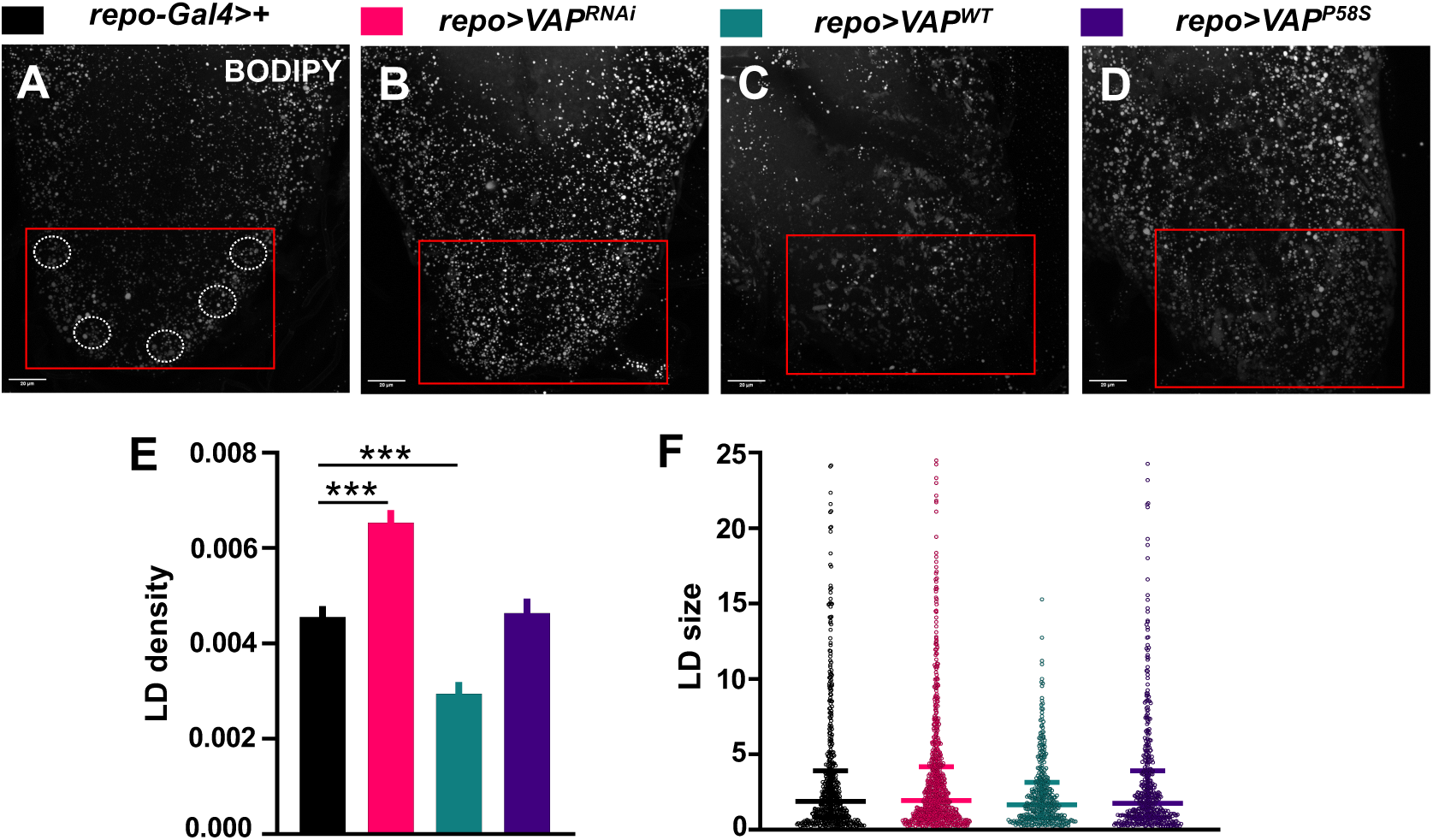
Glial *VAP* modulation affects LD homeostasis in the larval brain. **(A-D)** represent MIPs (20% Brightness) of the ventral nerve cord of the late 3^rd^ instar male larval brain stained with BODIPY. Scale bar = 20µm. Five ROIs are drawn at the tip of the VNC, outlined in red in the control (A), and used for LD analysis. *VAP* KD in glia shows an increase in LD number compared to control. Overexpression of *VAP^WT^*leads to a decrease in LD density, whereas *VAP^P58S^* OE shows no change compared to the control. **(E)** Quantitation of LD density (number of droplets per µm^3^) is plotted as Mean±SEM values in the graph. One-way ANOVA is used for mean comparison between genotypes, followed by Tukey’s test for multiple comparison correction, and p values are reported as *(<0.05), **(<0.01) and ***(<0.001). **(F)** LD size (volume in µm^3^) is plotted in the form of a scattered dot plot representing LDs present in ROIs of each genotype. The Kruskal-Wallis test was performed for mean rank comparison between genotypes, followed by Dunn’s test for multiple comparison correction. N=4-5 larval brains per genotype, n= 5 ROIs for each brain sample.

